# Integration of single-cell multiomic measurements across disease states with genetics identifies mechanisms of beta cell dysfunction in type 2 diabetes

**DOI:** 10.1101/2022.12.31.522386

**Authors:** Gaowei Wang, Joshua Chiou, Chun Zeng, Michael Miller, Ileana Matta, Jee Yun Han, Nikita Kadakia, Mei-Lin Okino, Elisha Beebe, Medhavi Mallick, Joan Camunas-Soler, Theodore dos Santos, Xiao-Qing Dai, Cara Ellis, Yan Hang, Seung K. Kim, Patrick E. MacDonald, Fouad R. Kandeel, Sebastian Preissl, Kyle J Gaulton, Maike Sander

**Affiliations:** Department of Pediatrics, University of California San Diego, La Jolla CA, USA; Pediatric Diabetes Research Center, University of California San Diego, La Jolla CA, USA; Biomedical Graduate Studies Program, University of California San Diego, La Jolla CA, USA; Center for Epigenomics, University of California San Diego, La Jolla CA, USA; Department of Bioengineering, Stanford University, Stanford, USA; Department of Pharmacology, University of Alberta, Edmonton, AB, Canada; Alberta Diabetes Institute, University of Alberta, Edmonton, AB, Canada; Department of Developmental Biology, Stanford University School of Medicine, Stanford, CA, USA; Departments of Medicine and of Pediatrics, Stanford University School of Medicine, Stanford, CA, USA; Stanford Diabetes Research Center, Stanford University School of Medicine, Stanford, CA, USA; Department of Clinical Diabetes, Endocrinology & Metabolism, City of Hope, Duarte, CA, USA; Institute of Experimental and Clinical Pharmacology and Toxicology, Faculty of Medicine, University of Freiburg, Freiburg, Germany; Institute for Genomic Medicine, University of California San Diego, La Jolla CA, USA; Department of Cellular and Molecular Medicine, University of California San Diego, La Jolla CA, USA; Max Delbrück Center for Molecular Medicine in the Helmholtz Association, Berlin, Germany

## Abstract

Altered function and gene regulation of pancreatic islet beta cells is a hallmark of type 2 diabetes (T2D), but a comprehensive understanding of mechanisms driving T2D is still missing. Here we integrate information from measurements of chromatin activity, gene expression and function in single beta cells with genetic association data to identify disease-causal gene regulatory changes in T2D. Using machine learning on chromatin accessibility data from 34 non-diabetic, pre-T2D and T2D donors, we robustly identify two transcriptionally and functionally distinct beta cell subtypes that undergo an abundance shift in T2D. Subtype-defining active chromatin is enriched for T2D risk variants, suggesting a causal contribution of subtype identity to T2D. Both subtypes exhibit activation of a stress-response transcriptional program and functional impairment in T2D, which is likely induced by the T2D-associated metabolic environment. Our findings demonstrate the power of multimodal single-cell measurements combined with machine learning for identifying mechanisms of complex diseases.

## Introduction

Pancreatic islets are comprised of multiple endocrine cell types with distinct functions in the regulation of glucose homeostasis and metabolism^1^. Islet endocrine cell types, in particular the insulin-producing beta cells, are known to exhibit substantial functional heterogeneity^2–4^. For example, in human islets, ∼20% of beta cells account for greater than 90% of the total insulin secreted at basal glucose levels^2^. Furthermore, gene expression studies at single-cell level have identified beta cell populations with distinct transcriptomic profiles^5^. Our group recently showed that beta cell subtypes can also be distinguished by chromatin activity in islets from non-diabetic (ND) donors^6^. Moreover, there is indication that beta cell subtypes could have relevance in type 2 diabetes (T2D), supported by the observation that subtypes defined by cell surface marker expression undergo an abundance shift in T2D^7^. How subtype-specific chromatin, transcriptomic and functional features relate to each and how changes in gene regulatory programs of beta cell subtypes could drive T2D pathogenesis is unknown.

T2D results from the interplay of both genetic and environmental factors. A change in beta cell function is a hallmark feature of pre-T2D^8, 9^, culminating in functional failure and eventual beta cell loss in T2D. To gain insight into mechanisms of beta cell failure in T2D, numerous studies have compared gene expression in islets from ND and T2D donors at both bulk^10, 11^ and single-cell level^5, 12–14^. However, these studies, including those at single-cell level, analyzed beta cells in aggregate, leaving unclear whether gene expression changes can be attributed to beta cell subtype shifts. Furthermore, it has been difficult to identify gene regulatory programs that are regulated in T2D across independent studies and cohorts, as evidenced by a meta-analysis^15^. Different islet procurement methods as well as heterogeneity due to confounding factors unrelated to disease impose analytical challenges of separating disease pathology from experimental noise. Given these limitations and challenges, insights into the gene regulatory changes causal to beta cell dysfunction in T2D will necessitate integration of information from single-cell measurements of chromatin activity, gene expression, and function with genetic association data, as well as analysis methods that minimize effects driven by disease-unrelated factors.

In this study, we measured chromatin activity and gene expression at single-cell level in a total of 34 islet preparations from ND, pre-T2D and T2D donors, using single nucleus ATAC-seq (snATAC-seq) and single nucleus RNA-seq (snRNA-seq). We developed a classifier based on machine learning from snATAC-seq data as an unbiased approach for identifying beta cell subtypes in heterogenous samples across disease. This approach identified two beta cell subtypes that change in abundance in T2D and can be reliably distinguished in data sets from independent cohorts. Using Patch-seq, which links cell electrophysiology as a proxy for insulin exocytosis to gene expression at single-cell level^16, 17^, we show that the two beta cell subtypes are functionally distinct in ND donors and impaired in function in T2D. Through gene regulatory network (GRN) analysis, we distinguish gene regulatory programs driving beta cell subtype identity from subtype-independent, T2D-associated changes. Finally, we describe the relationship of these gene regulatory programs to genetic risk of T2D which reveals a causal contribution of beta cell subtype identity to T2D pathogenesis.

## Results

### T2D affects chromatin state in beta cells

To map accessible chromatin in pancreatic islet cell types in healthy individuals and during T2D progression, we collected pancreatic islets from 11 ND, 8 pre-T2D and 15 T2D donors (34 total; **Supplementary Table 1a**) and profiled chromatin accessibility of individual cells by snATAC-seq (**Figure 1a**). After rigorous quality control (Methods and **Supplementary Figure 1a-g**), we annotated cell type identities based on chromatin accessibility at the promoter regions of known marker genes (**Figure 1b**, **Supplementary Figure 1h,i** and **Supplementary Table 1a,b**) and identified a total of 412,113 non-overlapping candidate *cis* regulatory elements (cCREs) (**Supplementary Table 2**).

**Figure 1.**
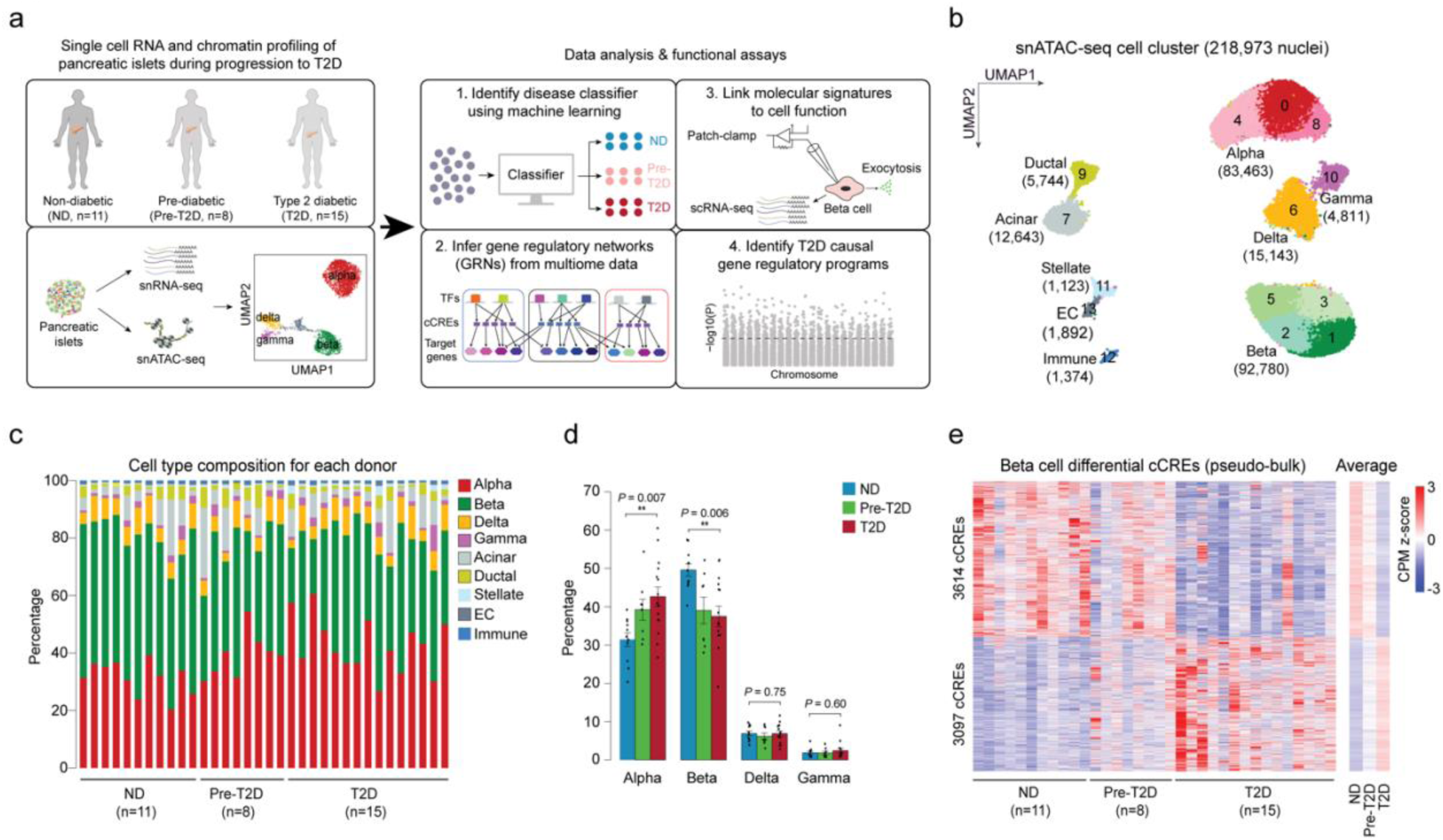
Beta cells exhibit changes in chromatin activity in type 2 diabetes. **(a)** Schematic outlining study design. snATAC-seq was performed on nuclei from pancreatic islets from 11 non-diabetic (ND), 8 pre-diabetic (pre-T2D) and 15 type 2 diabetic (T2D) human donors. Single nucleus multiome (ATAC+RNA) analysis was performed on a subset of donors (6 ND, 8 pre-T2D, 6 T2D). We used machine learning to identify classifiers for beta cells in ND, pre-T2D and T2D, inferred gene regulatory networks (GRNs), linked molecular signatures to beta cell function using Patch-seq, and identified T2D causal gene regulatory programs. **(b)** Clustering of chromatin accessibility profiles from 218,973 nuclei from non-diabetic, pre-diabetic, and T2D donor islets. Cells are plotted using the first two UMAP components. Clusters are assigned cell type identities based on promoter accessibility of known marker genes. The number of cells for each cell type cluster is shown in parentheses. EC, endothelial cells. **(c)** Relative abundance of each cell type based on UMAP annotation in Figure 1b. Each column represents cells from one donor. **(d)** Relative abundance of each islet endocrine cell type in ND, pre-T2D and T2D donor islets. Data are shown as mean ± S.E.M. (*n* = 11 ND, *n* = 8 pre-T2D, *n* = 15 T2D donors), dots denote data points from individual donors. \*\*\**P* < .001, \*\**P* < .01, \**P* < .05; ANOVA test with age, sex, BMI, and islet index as covariates. **(e)** Heatmap showing chromatin accessibility at cCREs with differential accessibility in beta cells from ND and T2D donors. Columns represent beta cells from each donor (ND, *n*=11; pre-diabetic, pre-T2D, *n*=8; T2D, *n*=15) and all ND, pre-T2D and T2D donors with accessibility of peaks normalized by CPM (counts per million).

Long-term T2D leads to beta cell loss^18^, and therefore we assessed changes in cell type composition between islets from ND, pre-T2D and T2D donors. Cell type composition exhibited substantial donor heterogeneity (**Figure 1c**), consistent with previous reports^11^. Relative beta cell numbers were significantly reduced in T2D compared to ND donor islets (*P*=0.006, ANOVA test), whereas relative alpha cell numbers were increased (*P*=0.007, ANOVA test; **Figure 1d**). By contrast, relative delta or gamma cell numbers were similar between ND and disease groups (**Figure 1d**).

Characterization of cell type-resolved changes in chromatin accessibility during T2D progression can reveal gene regulatory mechanisms leading to T2D. Considering biological (age, sex, BMI) and technical (islet index, fraction of reads overlapping TSS, total read counts) covariates (Methods and **Supplementary Figure 2**), we identified cCREs with differential activity between ND, pre-T2D and T2D donors in aggregate beta cells (“pseudo-bulk”)^19^. We observed substantial differences in beta cell chromatin activity between ND and T2D donors, where 3,097 and 3,614 cCREs gained and lost accessibility in T2D, respectively (FDR<0.1, p-values adjusted with the Benjamini-Hochberg method; **Figure 1e** and **Supplementary Table 3a**). Of the 6,711 differential cCREs in our cohort, 78.8% (5,291/6,711) showed consistent changes in an independent cohort of ND (*n*=15) and T2D donors (*n*=5) (*P*<2.2×10^-16^, Binominal test; **Supplementary Figure 3**; see Methods for data source), demonstrating robustness of our findings. There were no beta cell differential cCREs between ND and pre-T2D donors, and only a few between pre-T2D and T2D donors (**Supplementary Table 3b**). The same result was obtained after down-sampling to match donor numbers in the ND, pre-T2D and T2D groups (**Supplementary Table 3c-h**). To explore whether intermediate chromatin activity changes were present in pre-T2D samples, we calculated the percentage of T2D versus ND differential beta cell cCREs exhibiting directionally concordant changes in pre-T2D. We found that 96% of cCREs gaining (2975/3097; *P*<2.2×10^-16^, Binominal test) and 97% of cCREs losing (3614/3614; *P*<2.2×10^-16^, Binominal test) accessibility in T2D exhibited directionally concordant changes in pre-T2D and T2D (**Figure 1e**). Therefore, although T2D-relevant changes in beta cell chromatin activity are subtly present in pre-T2D, chromatin activity more closely resembles ND than T2D donors.

To identify potential effects of T2D on chromatin in non-beta islet cell types, we tested cCREs for differential activity in alpha, delta and gamma cells, but found no or very few regulated cCREs (13 differential alpha cell cCREs between ND and T2D donors; **Supplementary Table 4**). We next sought to determine whether this is due to a lack of power as a result of lower cell numbers, and therefore down-sampled beta cell numbers to more closely match the numbers of alpha and delta cells. Down-sampling to 15,000 beta cells identified 1,070 differential cCREs, whereas no differential cCREs were identified in similar numbers of alpha and delta cells (FDR<0.1; **Supplementary Figure 4**). To confirm disease-specificity of the identified beta cell differential cCREs, we further called differential cCREs after shuffling the disease status of donors (FDR<0.1, p-values adjusted with the Benjamini-Hochberg method); however, we identified no differential cCREs in either beta or alpha cells. This analysis supports the conclusion that effects of T2D on chromatin accessibility are more subtle in non-beta islet cell types compared to beta cells.

### Machine learning identifies two beta cell subtypes based on chromatin activity

The T2D-associated chromatin activity changes in aggregate beta cells (**Figure 1e**) could be due to a shift in beta cell subpopulations, a shift in chromatin activity in individual beta cells, or both (**Figure 2a**). To distinguish between these possibilities, we first re-clustered beta cells and identified three beta cell clusters (**Supplementary Figure 5a**). However, none of the clusters showed a preference for beta cells from pre-T2D or T2D donors (**Supplementary Figure 5b**). A shortcoming of clustering and dimensionality reduction is that factors unrelated to disease can drive subtype identity in the clustering and obscure disease-relevant shifts. To circumvent these limitations, we applied machine learning^20^ (see Methods) by training a classifier on individual beta cells and testing its ability to distinguish beta cell chromatin profiles from ND, pre-T2D and T2D donors (∼240k cCREs across all 34 donors). To eliminate donor-specific effects during model training and to test whether beta cells from ND, pre-T2D and T2D donors can be distinguished, we removed beta cells from one donor at a time in the testing group while using remaining donors as a training group. To determine the accuracy of the classifier for predicting disease state, we compared predictions of the classifier to the annotated disease state for each donor. If chromatin activity changes gradually in individual beta cells during progression from the ND to the pre-T2D and T2D state (Scenario 2, **Supplementary Figure 5c**), the classifier should exhibit high prediction accuracy in all three states. By contrast, if T2D progression is associated with a shift in beta cell subtypes (Scenario 3, **Supplementary Figure 5c**), prediction accuracy will depend on the prevalence of the dominant beta cell subtype. The classifier predicted beta cells from ND and T2D donors with ∼60% accuracy, while the prediction accuracy of beta cells from pre-T2D donors was only at ∼5% (**Supplementary Figure 5d,e**). This indicates presence of two major beta cell subtypes, one being enriched in ND donors and the other being enriched in donors with T2D (Scenario 3). Confirming this conclusion, similar prediction accuracies (ND = 48.4%, pre-T2D = 16.2%, T2D = 49.0%) were observed after down-sampling beta cells from ND and T2D donors to numbers from pre-T2D donors.

**Figure 2.**
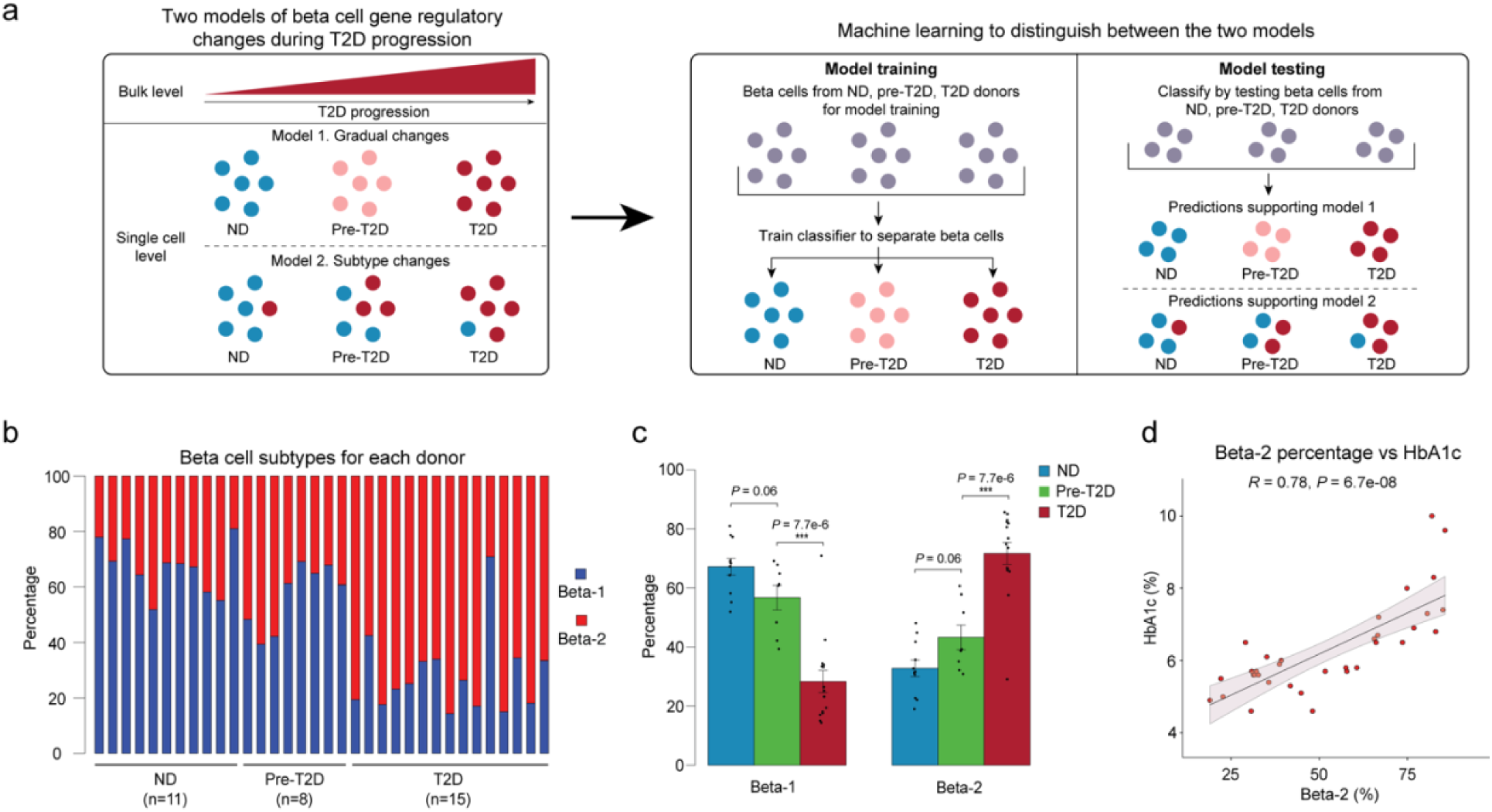
Machine learning identifies two beta cell subtypes with differential abundance in T2D. **(a)** Schematic outlining the machine learning-based approach to distinguish two models that could account for gene regulatory changes in beta cells in T2D. **(b)** Relative abundance of beta-1 and beta-2 cells identified by machine learning. Each column represents cells from one donor. **(c)** Relative abundance of each beta cell subtype in ND, pre-T2D and T2D donor islets. Data are shown as mean ± S.E.M. (*n* = 11 ND, *n* = 8 pre-T2D, *n* = 15 T2D donors), dots denote data points from individual donors. \*\*\**P* < .001; ANOVA test with age, sex, BMI, and islet index as covariates. **(d)** Pearson correlation between relative abundance of beta-2 cells and HbA1c across donors (*n* = 11 ND, *n* = 8 pre-T2D, *n* = 15 T2D donors).

The same analysis for alpha and delta cells showed prediction accuracies for ND, pre-T2D and T2D of 20-30% (**Supplementary Figure 5f-i**), which is close to randomness (Scenario 1, **Supplementary Figure 5c**). This suggests that alpha and delta cells from ND, pre-T2D and T2D donors are indistinguishable, agreeing with the finding that there were no differentially active cCREs.

By applying reiterative training and testing steps on beta cells from only ND and T2D donors (Methods, **Supplementary Figure 5j**), we next established a classifier capable of distinguishing the beta cell subtype enriched in ND donors (hereafter beta-1) and T2D donors (hereafter beta-2; **Supplementary Table 5**) and calculated their relative abundance in each donor (**Figure 2b**). Beta-1 cells were less abundant in T2D donors (28.3±3.7% of beta cells) compared to ND donors (67.2±2.8% of beta cells), whereas beta-2 cells were more abundant in T2D donors (32.7±2.8% beta-2 in ND and 71.7±3.8% beta-2 in T2D; **Figure 2c**). There was a small, non-significant decrease in beta-1 and increase in beta-2 cells in pre-T2D compared to ND donors (**Figure 2c**), suggesting that the subtype shift mostly occurs in T2D. At the level of individual donors, the abundance of beta-2 cells positively correlated with HbA1c (**Figure 2d**), which is an index for long-term glycemic control. The percentage of beta-2 cells was unrelated to sex, BMI, or the islet index as a technical confounding factor, but showed a nominal but small positive correlation with age (**Supplementary Figure 6a-d**).

To further confirm that beta cell subtype identity shifts in T2D, we validated our findings using independent data sets and analysis methods. Testing our classifier on snATAC-seq data from another cohort (15 ND and 5 T2D; data source see Methods) revealed similar proportions of beta-1 and beta-2 cells in ND and T2D donors as observed in our cohort (**Supplementary Figure 6e,f**). As in our cohort, the abundance of beta-1 cells decreased and beta-2 cells increased in T2D, showing robustness of our classifier for identifying beta cell subtypes and T2D-associated changes. Next, we tested whether methods other than machine learning can confirm the presence of the two beta cell subtypes. Since the machine learning approach identified the subtype shift as the most prominent gene regulatory change in T2D, we predicted that many of the differentially active cCREs in aggregate beta cells from ND and T2D donors (see **Figure 1e**) represent subtype-specific cCREs. To test this, we clustered beta cells based on cCREs with differential activity in aggregate beta cells from T2D donors. Indeed, this clustering identified two beta cell populations with differential abundance in T2D (**Supplementary Figure 6g-k**). Importantly, beta cells in cluster 1 and cluster 2, respectively, overlapped significantly with beta-1 and beta-2 cells identified by machine learning (*P* < 2.2e-16, exact binomial test; **Supplementary Figure 6l**), showing robustness of subtype identity assignments across methods.

### The two beta cell subtypes are transcriptionally and functionally distinct

To understand the gene expression programs that distinguish the two beta cell subtypes, we profiled gene expression and chromatin accessibility jointly from the same nuclei (Single-cell Multiome, 10x Genomics) in a subset of donors (6 ND, 8 pre-T2D, 6 T2D; **Supplementary Table 1a**). We (i) isolated beta cells by independently clustering based on snATAC-seq and snRNA-seq data (**Supplementary Figure 7a,b**), (ii) showed that clustering beta cells based on genes linked to cCREs with differential activity in T2D (see **Figure 1e**) separates beta-1 and beta-2 subtypes defined by snATAC-seq (**Supplementary Figure 7c,d**) and (iii) identified differential cCREs (*n*=34 donors) and differentially expressed genes (*n*=20 donors) between beta-1 and beta-2 cells (Methods and **Figure 3a,b**). Changes in distal and promoter cCRE activity positively correlated with changes in gene expression (Methods and **Supplementary Figure 7e,f**). Genes with higher expression and chromatin accessibility in beta-2 compared to beta-1 cells included insulin (*INS*) and positive regulators of insulin secretion, such as synaptotagmin 1 (*SYT1*) and glucokinase (*GCK*), as well as the transcription factor (TF) *PAX6* which positively regulates insulin gene transcription^21^ (**Figure 3b,c**, **Supplementary Figure 7g** and **Supplementary Table 6**). Beta-1 cells expressed higher levels of the TFs *HNF1A* and *HNF4A* (**Figure 3b,c** and **Supplementary Table 6**). Accordingly, HNF1A and HNF4A motifs were enriched at cCREs with higher activity in beta-1 than beta-2 cells, while NEUROD1, E2A and NF1 motifs were enriched at cCREs more active in beta-2 cells (**Figure 3d** and **Supplementary Table 7**). Together, this analysis identifies concordant gene regulatory and transcriptomic features that distinguish the two beta cell subtypes. We further validated the beta cell subtypes using human islet scRNA-seq data from three independent cohorts^5, 12, 22^ (Methods). In each cohort, clustering of beta cells based on beta-1 versus beta-2 differentially expressed genes identified two beta cell populations (**Supplementary Figure 8a,d,g**) with directionally similar gene expression differences as in beta-1 versus beta-2 cells (**Supplementary Figure 8b,e,h**). Furthermore, the relative abundance of beta-1 and beta-2 cells in ND and T2D donors was consistent with the observations in our cohort (**Supplementary Figure 8c,f,i**).

**Figure 3.**
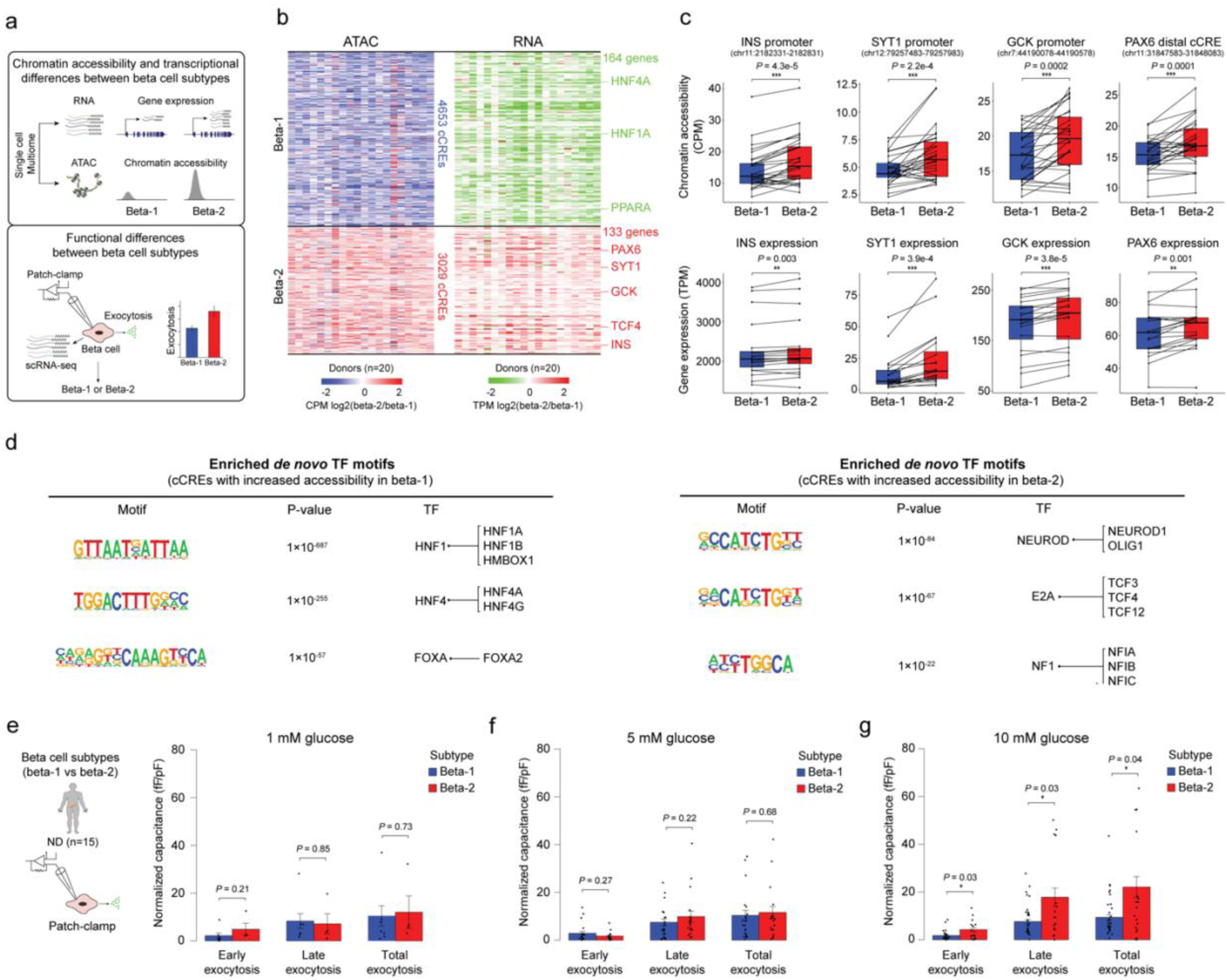
The two beta cell subtypes are distinguished by chromatin activity, gene expression and function. **(a)** Workflow to link beta cell subtype chromatin activity to gene expression using islet single nucleus multiome (ATAC+RNA) data and gene expression to function using Patch-seq. **(b)** Heatmap showing log_2_ differences (beta-2/beta-1) in chromatin accessibility at cCREs with differential accessibility between beta cell subtypes (left, paired t-test, FDR < 0.05, *P*-values adjusted with the Benjamini-Hochberg method) and log_2_ differences (beta-2/beta-1) in gene expression of cCRE target genes with differential expression between beta cell subtypes (right, paired t-test, FDR < 0.15, *P*-values adjusted with the Benjamini-Hochberg method). Rows represent differential cCREs or genes, columns represent donors (total 20, ND, *n*=6; pre-T2D, *n*=8; T2D, *n*=6). Representative genes are highlighted. Accessibility of cCREs is normalized by CPM (counts per million) and gene expression by TPM (transcripts per million). **(c)** Bar plots showing cCRE accessibility (top) and gene expression (bottom) of representative genes in beta-1 and beta-2 cells. Proximal region of *INS* (chr11:2182331-2182831), *SYT1* (chr12:79257483-79257983), *GCK* (chr7:44190078-44190578), *PAX6* (chr11:31847583-31848083). Accessibility of peaks is normalized by CPM and gene expression by TPM. Paired t-test. **(d)** Transcription factor (TF) motif enrichment at cCREs with higher accessibility in beta-1 compared to beta-2 cells (left) or higher accessibility in beta-2 compared to beta-1 cells (right) against a background of all cCREs in beta cells using HOMER. The top three enriched *de novo* motifs, their *P*-values, and best matched known TF motif are shown. **(e)** Bar plots from Patch-seq analysis showing early, late and total exocytosis in beta-1 (10 cells from 4 ND donors) and beta-2 cells (4 cells from 4 ND donors) stimulated with 1 mM glucose. Data are shown as mean ± S.E.M., dots denote data points from individual cells. ANOVA test with age, sex, and BMI as covariates. **(f)** Bar plots from Patch-seq analysis showing early, late and total exocytosis in beta-1 (26 cells from 10 ND donors) and beta-2 cells (20 cells from 9 ND donors) stimulated with 5 mM glucose. ANOVA test with age, sex, and BMI as covariates. **(g)** Bar plots from Patch-seq analysis showing early, late and total exocytosis in beta-1 (42 cells from 5 ND donors) and beta-2 cells (23 cells from 6 ND donors) stimulated with 10 mM glucose. \**P* < .05, ANOVA test with age, sex, and BMI as covariates.

The higher expression of insulin and genes associated with insulin secretion in beta-2 cells indicates possible functional differences between the beta cell subtypes. To test this, we leveraged Patch-seq (electrophysiological measurements + scRNA-seq) data from human islets (15 ND donors; **Figure 3a**) in which we confirmed the two beta cell subtypes (**Supplementary Figure 8j-l**). Comparison of exocytosis in beta-1 and beta-2 cells from ND donors revealed higher exocytosis in beta-2 than beta-1 cells in high glucose (10 mM) (**Figure 3e-g**). This finding suggests that beta-2 cells, which is the minority population in ND donors (**Figure 2c**), release more insulin in response to glucose than beta-1 cells. In sum, these results demonstrate that a classifier based on machine learning of epigenomic profiles can discern beta cell subtypes with distinct transcriptomic and functional features. The less abundant beta cell subtype in ND donors expresses insulin and exocytotic genes at higher levels and exhibits increased exocytosis under high glucose conditions in ND donors.

### A bistable transcriptional circuit maintains the two beta cell subtypes

The presence of two distinct beta cell subtypes raises the question of how the two beta cell states are maintained. To uncover transcriptional mechanisms of beta subtype maintenance, we inferred beta cell GRNs, linking TFs to cCREs and their target genes (Methods and **Figure 4a**). Briefly, we performed TF binding motif analysis at beta cell cCREs, focused on TFs expressed in beta cells, linked cCREs to genes based on proximity and co-accessibility, and calculated the correlation between TF and gene expression in aggregate beta-1 and beta-2 cells for each donor from our multiome data (*n*=20 donors; Methods). For each TF (total of 266 TFs) we identified target genes with positive or negative expression correlation with the TF (**Supplementary Table 8**). The positively and negatively regulated TF-gene modules comprised a median number of 600 and 505 target genes, respectively.

**Figure 4.**
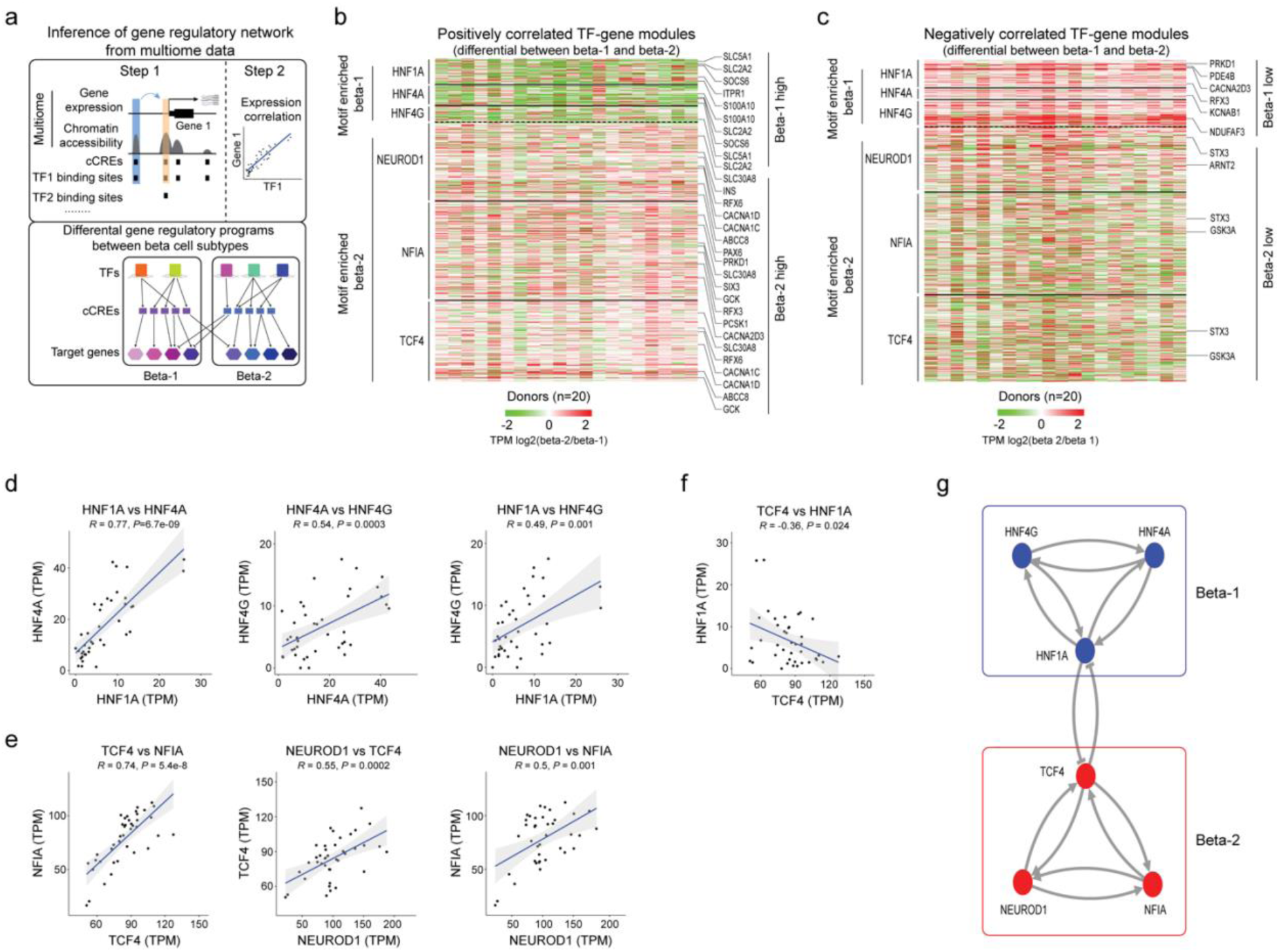
Gene regulatory networks defining the two beta cell subtypes. **(a)** Schematic outlining the inference of beta cell gene regulatory networks and differential gene regulatory programs (TF-gene modules) between beta cell subtypes. **(b)** Heatmap showing log_2_ differences (beta-2/beta-1) in expression for genes positively regulated by TFs (HNF1A, HNF4A, HNF4G) with higher activity in beta-1 compared to beta-2 cells and TFs (NEUROD1, NFIA and TCF4) with higher activity in beta-2 compared to beta-1 cells (see Methods). Representative target genes of individual TFs are highlighted. Gene expression is normalized by TPM (transcripts per million). **(c)** Heatmap showing log_2_ differences (beta-2/beta-1) in expression for genes negatively regulated by TFs (HNF1A, HNF4A, HNF4G) with higher activity in beta-1 compared to beta-2 cells and TFs (NEUROD1, NFIA, TCF4) with higher activity in beta-2 compared to beta-1 cells (see Methods). Representative target genes of individual TFs are highlighted. Gene expression is normalized by TPM (transcripts per million). **(d, e, f)** Pearson correlation of expression levels between indicated TFs across pseudo-bulk RNA profiles from each beta cell subtype (40 dots in total: 20 donors including *n* = 6 ND, *n* = 8 pre-T2D, *n* = 6 T2D). **(g)** A bistable circuit established by positive feedback between HNF1A, HNF4A and HNF4G, positive feedback between NEUROD1, NFIA and TCF4, and mutual repression between HNF1A and TCF4.

Next, we sought to isolate TF-gene modules with differential regulation between beta-1 and beta-2 cells. First, we conducted gene set analysis (GSA)^23–25^ to identify modules where genes exhibit a significant difference in expression between beta-1 and beta-2 cells in both the positively and negatively regulated module for a given TF (*P*<0.05; Methods). Second, we filtered TF-gene modules based on TF motifs enriched at cCREs with differential activity between beta-1 and beta-2 cells (see **Figure 3d** and **Supplementary Table 7**). This analysis revealed gene modules positively and negatively regulated by HNF1A, HNF4A and HNF4G with higher and lower expression, respectively, in beta-1 than beta-2 cells; and, conversely, gene modules positively and negatively regulated by NEUROD1, NFIA and TCF4 with higher and lower expression, respectively, in beta-2 than beta-1 cells (**Figure 4b,c**). Among the genes positively regulated by HNF1A, HNF4A and HNF4G were known regulators of insulin secretion, including the glucose transporter *SLC2A2*^26^, the suppressor of cytokine signaling *SOCS6*^27^, the calcium binding protein *S100A10*^28^, and the ligand-gated calcium channel *ITPR1*^29^ (**Figure 4b**, **Supplementary Figure 9a** and **Supplementary Table 8**). Likewise, positively regulated targets of NEUROD1, NFIA and TCF4 included many genes with established roles in beta cell function (*SLC30A8, RFX6, ABCC8, INS, GCK, PCSK1*) (**Figure 4b**, **Supplementary Figure 9b** and **Supplementary Table 8**).

To identify mechanisms that reinforce beta cell subtype identity, we analyzed how the beta-1 and beta-2 subtype-defining TFs are regulated. For *HNF1A, HNF4A* and *HNF4G*, promoter chromatin accessibility and expression were higher in beta-1 than beta-2 cells (**Supplementary Figure 9c,d**). Conversely, *TCF4* and *NFIA* exhibited higher promoter accessibility and expression in beta-2 cells (**Supplementary Figure 9e,f**). The beta cell subtype enrichment of each one of these TFs appears to be reinforced by auto-regulatory and cross-regulatory feedback loops. For example, we found beta-1 versus beta-2 differentially active cCREs at *HNF1A*, *HNF4A* and *HNF4G* containing predicted binding sites for HNF1A, HNF4A and HNF4G (**Supplementary Figure 9g**) and observed positive correlation in expression between these TFs in beta cells across donors (**Figure 4d**). Similar positive feedback loops were identified between NEUROD1, NFIA and TCF4 (**Supplementary Figure 9h** and **Figure 4e**). HNF1A and TCF4 showed negative feedback (**Supplementary Figure 9i** and **Figure 4f**), suggesting that beta cell subtype identity is maintained by a bistable transcriptional switch between HNF1A and TCF4 which is reinforced by positive feedback loops between beta subtype-defining TFs (**Figure 4g**). Together, this analysis identifies a core network of TFs and their target genes governing beta cell subtype identity.

### T2D-related functional and gene regulatory changes in beta cells

Beta-2 cells exhibit higher insulin exocytosis than beta-1 cells in ND donors (**Figure 3e-g**); however, beta-2 cells increase in abundance in T2D (**Figure 2c**). These observations are difficult to reconcile with the T2D-associated decline in beta cell function^8, 16^. To determine whether beta-1 and/or beta-2 cells undergo functional change during T2D progression, we compared insulin exocytosis in beta-1 and beta-2 cells from ND (*n*=15), pre-T2D (*n*=16), and T2D (*n*=14) donors using Patch-seq. There was no difference in exocytosis at stimulatory glucose (5 mM and 10 mM) between ND and pre-T2D donors in either beta-1 and beta-2 cells. By contrast, both beta-1 and beta-2 cells exhibited decreased exocytosis in T2D compared to pre-T2D donors (**Figure 5a,b**). Thus, both beta-1 and beta-2 cells exhibit functional impairment in T2D, consistent with an overall decline in beta cell function in T2D.

**Figure 5.**
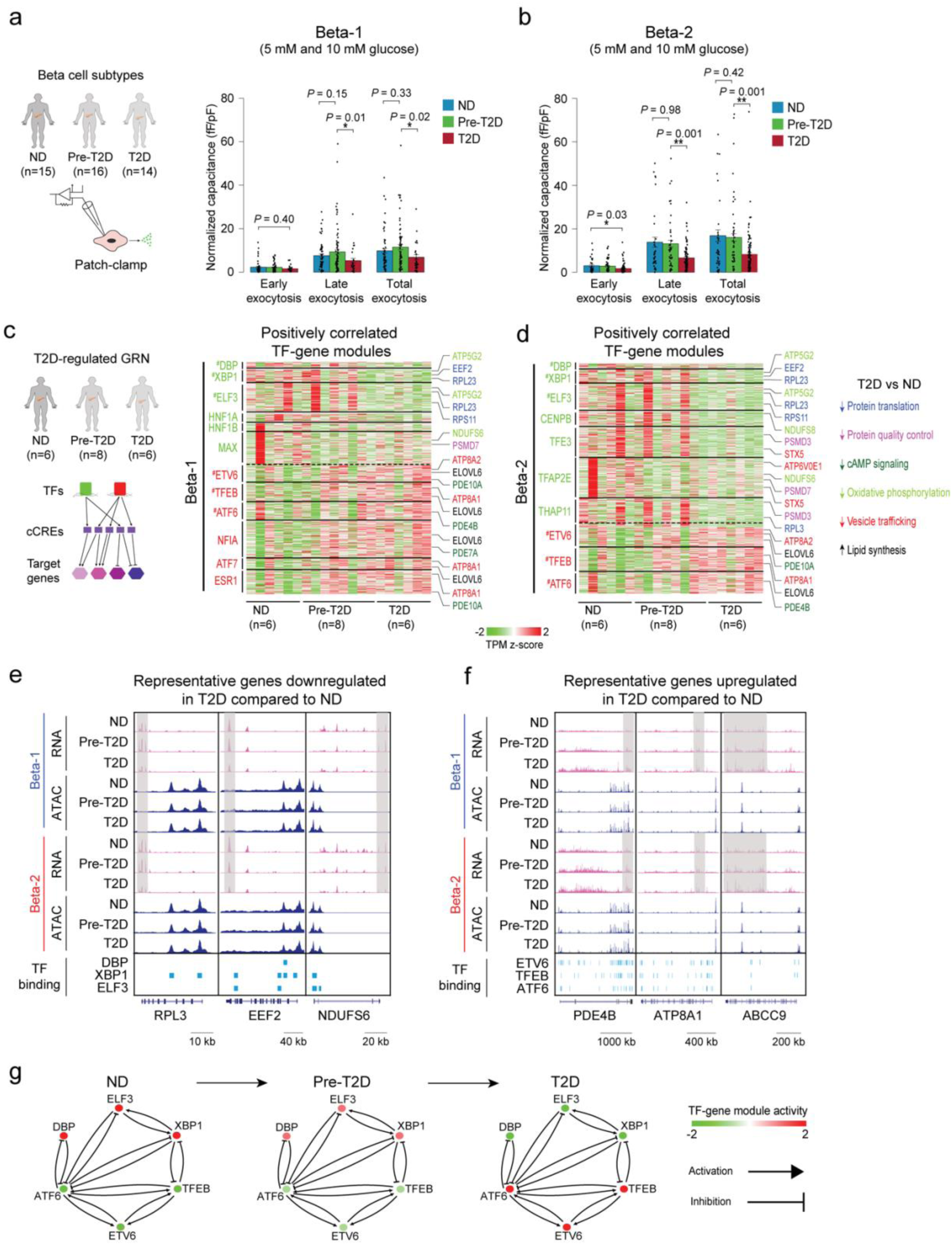
Beta cell functional and gene regulatory changes in T2D. **(a)** Bar plots from Patch-seq analysis showing early, late and total exocytosis in beta-1 cells from ND (68 cells from 11 donors), pre-T2D (91 cells from 14 donors) and T2D donors (35 cells from 7 donors) stimulated with 5 mM or 10 mM glucose. Data are shown as mean ± S.E.M., dots denote data points from individual cells. \**P* < .05, ANOVA test with age, sex, and BMI as covariates. **(b)** Bar plots from Patch-seq analysis showing early, late and total exocytosis in beta-2 cells from ND (43 cells from 10 donors), pre-T2D (57 cells from 14 donors) and T2D donors (131 cells from 14 donors) stimulated with 5 mM or 10 mM glucose. \**P* < .05, \*\**P* < .01, ANOVA test with age, sex, and BMI as covariates. **(c)** Heatmap showing expression of genes positively regulated by TFs (green) with higher activity in ND compared to T2D beta-1 cells (see Methods) and TFs (red) with lower activity in ND compared to T2D beta-1 cells (*n*=6 ND, *n*=8 pre-T2D, *n*=6 T2D donors). Representative target genes of individual TFs are highlighted and classified by biological processes. Gene expression is normalized by TPM (transcripts per million). ^#^ denotes TFs with decreased or increased expression in T2D in both beta-1 and beta-2 cells. **(d)** Heatmap showing expression of genes positively regulated by TFs (green) with higher activity in ND compared to T2D beta-2 cells (see Methods) and TFs (red) with lower activity in ND compared to T2D beta-2 cells (*n*=6 ND, *n*=8 pre-T2D, *n*=6 T2D donors). Representative target genes of individual TFs are highlighted and classified by biological processes. Gene expression is normalized by TPM (transcripts per million). ^#^ denotes TFs with decreased or increased expression in T2D in both beta-1 and beta-2 cells. **(e)** Genome browser tracks showing aggregate RNA and ATAC read density at representative genes (*RPL3*, *EEF2*, *NDUFS6*) downregulated in T2D relative to ND for both beta-1 and beta-2 cells. Downregulated regions in T2D beta cells are indicated by grey shaded boxes. Beta cell cCREs with binding sites for downregulated TFs in both beta-1 and beta-2 cells (DBP, XBP1, ELF3) are shown. All tracks are scaled to uniform 1×10^6^ read depth. **(f)** Genome browser tracks showing aggregate RNA and ATAC read density at representative genes (*PDE4B*, *ATP8A1*, *ABCC9*) upregulated in T2D relative to ND for both beta-1 and beta-2 cells. Upregulated regions in T2D beta cells are indicated by grey shaded boxes. Beta cell cCREs with binding sites for upregulated TFs in both beta-1 and beta-2 cells (ETV6, TFEB, ATF6) are shown. All tracks are scaled to uniform 1×10^6^ read depth. **(g)** Cross regulation between TFs with activity change in T2D in both beta cell subtypes (from Figure 5c,d). The color code for TFs in ND, pre-T2D and T2D donors reflects their expression change during T2D progression.

To understand the molecular basis of these functional changes in beta cells in T2D, we analyzed T2D-associated alterations in gene regulatory programs within beta-1 and beta-2 cell populations. To this end, we identified differentially active cCREs in beta-1 and beta-2 cells between ND, pre-T2D and T2D donors (Methods and **Supplementary Table 9**). Both beta-1 and beta-2 cells exhibited significant changes in chromatin activity between ND and T2D donors (**Supplementary Figure 10a,b)**. Consistent with the findings in aggregate beta cells (**Figure 1e**), there were few differential cCREs between ND and pre-T2D or pre-T2D and T2D donors (**Supplementary Table 9**). However, both beta cell subtypes showed subtle changes in chromatin activity in pre-T2D that were directionally concordant with T2D-associated changes (beta-1 and beta-2: 99% and 98% of cCREs losing and gaining activity, respectively; *P*<2.2×10^-16^, Binominal test).

To further characterize T2D-induced gene regulatory changes in each beta cell subtype, we inferred T2D-regulated GRNs by identifying TF-gene modules in beta-1 and beta-2 cells with changes in T2D (Methods; **Figure 5c,d**, **Supplementary Figure 10c,d** and **Supplementary Table 10**). Consistent with our analysis of chromatin accessibility, there were no modules with differential regulation between ND and pre-T2D. The analysis revealed TFs that regulate gene modules in both beta cell subtypes as well as TFs regulating gene modules in only one subtype in T2D. TFs driving T2D-associated gene expression changes in both subtypes included the signal-dependent TFs DBP, ELF3, XBP1, TFEB, ETV6, and ATF6 (**Figure 5c,d**). These TFs are regulated by cell extrinsic stimuli including nutrients and circadian cues and are known mediators of the cellular stress response^30–32^. This suggests that T2D-associated changes in the extracellular environment, such as elevated glucose, affect gene expression in both beta cell subtypes. Interestingly, we observed regulation of HNF1A- and NFIA-driven gene modules in T2D in beta-1 but not beta-2 cells (**Figure 5c,d** and **Supplementary Figure 10c,d**). Down-regulation of the HNF1A module and up-regulation of the NFIA module in beta-1 cells indicates that beta-1 cells shift towards beta-2 identity in T2D, in accordance with the T2D-associated decrease in beta-1 cell abundance (**Figure 2c**).

Processes associated with genes regulated in both beta cell subtypes in T2D included protein translation and protein quality control, cAMP signaling, oxidative phosphorylation, vesicle trafficking, and lipid metabolism (**Figure 5c,d**, **Supplementary Figure 10c,d** and **Supplementary Table 11**). These processes are known to be affected by the stress response in beta cells and to alter beta cell function^33^, consistent with the functional changes of beta cells in T2D. For example, downregulated modules in T2D included genes encoding mitochondrial electron transport chain proteins (*NDUFS6, NDUFS8*, *ATP5G2*), syntaxins (*STX5*), and multiple ribosomal proteins important for protein translation (*RPL3, EEF2, EIF3I,*) (**Figure 5c-e** and **Supplementary Figure 10c,d**). These gene expression changes are predicted to reduce insulin production and secretion. By contrast, gene modules with increased expression in T2D included negative regulators of cAMP signaling (*PDE4B, PDE7A*) (**Figure 5c,d,f** and **Supplementary Figure 10c,d**), known to dampen glucose-stimulated insulin secretion^34^. Furthermore, we observed upregulation of regulators of insulin secretion including K_ATP_ channel subunits (*ABCC9*)^35^ and P4-ATPases (*ATP8A1*, *ATP8A2*)^36^ as well as lipogenic enzymes (*ELOVL6, ELOVL7*) which module the endoplasmic reticulum (ER) stress response^37^ and inhibit insulin secretion^38^ (**Figure 5c,d,f** and **Supplementary Figure 10c,d**). Of interest, distinct TFs regulated similar genes in the two beta cell subtypes, exemplified by MAX regulating *RPL5*, *NDUFS6, PDE7B,* and *ELOVL6* in beta-1 cells and TFAP2E regulating the same genes in beta-2 cells. Thus, our analysis identifies a core gene regulatory program comprised of signal-dependent TFs associated with the stress response that converge on similar genes that are dysregulated in T2D.

To understand the gene regulatory mechanisms leading to functional changes in T2D, we defined the regulatory relationship between the TFs driving gene expression changes in both beta cell subtypes. We observed positive correlation in expression across donors among TFs downregulated (*XBP1, ELF3*) and upregulated (*ETV6, TFEB, ATF6*) in T2D, respectively (**Supplementary Figure 10e,f**), as well as negative correlation between TFs changing in opposite directions in T2D (**Supplementary Figure 10g**). This suggests that positive and negative feedback loops between these TFs reinforce T2D-related gene expression changes. Donor-specific quantification of gene activity in each TF-gene module across disease states revealed subtle changes between ND and pre-T2D donors and more pronounced changes between pre-T2D and T2D (**Figure 5g**), consistent with observed patterns of chromatin activity.

### Genetic risk of T2D affects beta cell subtype regulation

Hundreds of genetic risk loci have been identified for T2D, many of which impact beta cell function^39^. We thus leveraged the highly polygenic inheritance of T2D to determine the beta cell transcriptional programs that contribute to T2D risk. We tested for enrichment of fine-mapped T2D risk variants in cCREs with increased activity in the beta-1 against the beta-2 subtype and vice versa compared to a background of permuted cCREs derived from all beta cell cCREs. We observed strong enrichment of T2D risk variants in cCREs with increased activity for both beta-1 and beta-2 subtypes compared to background cCREs (beta-1 logOR=1.33, *P*=1.8×10^-3^; beta-2 logOR=1.75, *P*=1.5×10^-4^; **Figure 6a**). Next, we tested for enrichment of fine-mapped T2D risk variants in cCREs with increased or decreased activity in beta-1 and beta-2 subtypes across the T2D disease state. We did not observe significant enrichment of these cCREs for T2D risk variants, although there was nominal evidence (*P*<.05) for enrichment of beta-2 cCREs with higher activity in T2D (**Supplementary Figure 11**).

**Figure 6.**
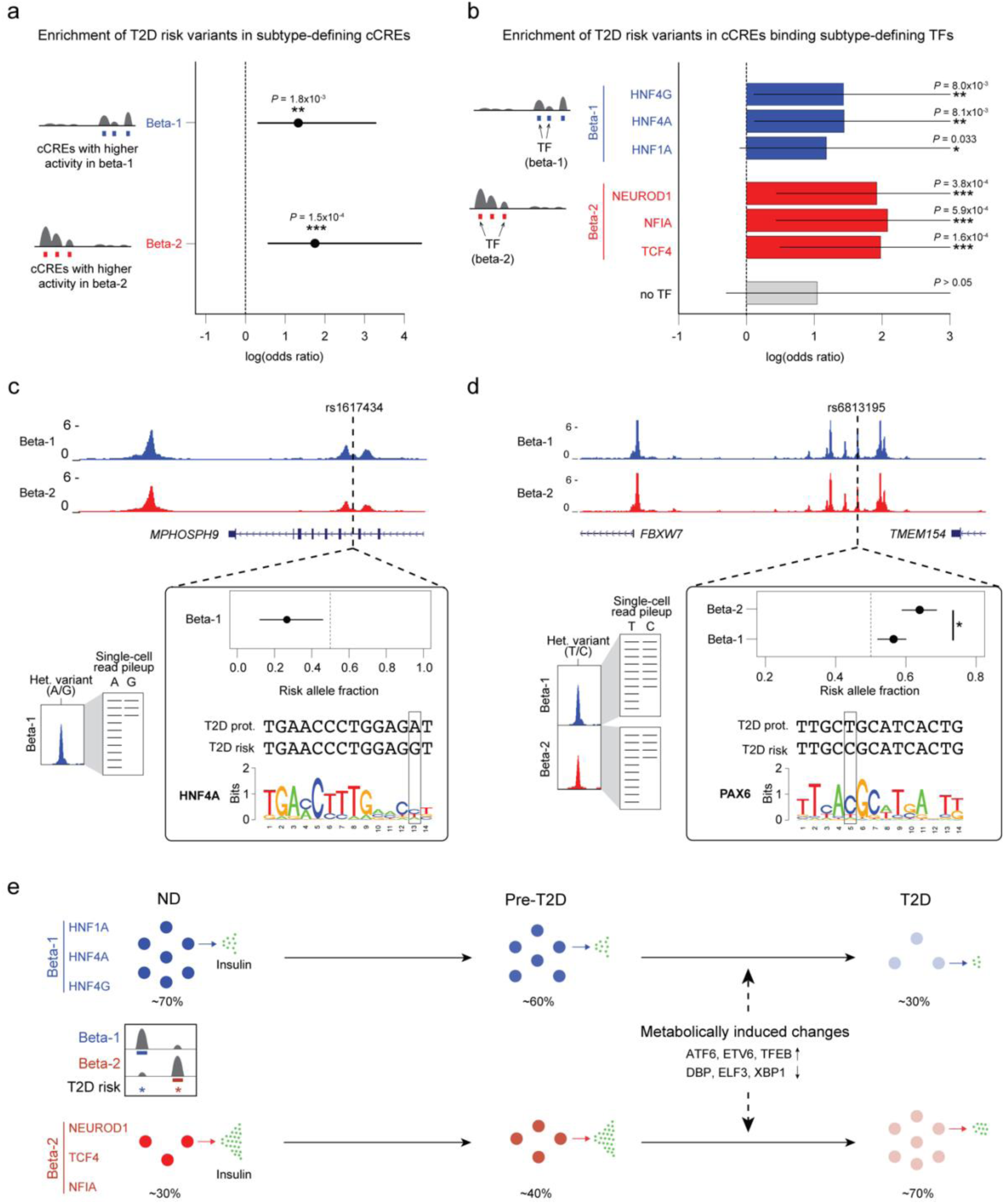
T2D risk variants affect beta cell subtype chromatin activity. **(a)** Enrichment of fine-mapped T2D risk variants for cCREs defining the beta-1 and beta-2 subtype. Values represent log odds ratios and 95% confidence intervals. **(b)** Enrichment of fine-mapped T2D risk variants for cCREs defining the beta-1 and beta-2 subtype bound by each TF, or not bound by any of the listed TFs (‘no TF’). Values represent log odds ratios and 95% confidence intervals. **(c)** Fine-mapped T2D risk variant rs1617434 at the *MPHOSPH9* locus overlaps a cCRE defining the beta-1 subtype. The T2D risk allele of this variant decreases beta-1 chromatin accessibility and disrupts a predicted binding site for HNF4A. The values for allelic imbalance represent the fraction of reads from the risk allele and the 95% confidence interval. On the left is a schematic describing allelic imbalance mapping in reads from the beta-1 subtype. **(d)** Fine-mapped T2D risk variant rs6813185 at the *TMEM154/FBXW7* locus overlaps a cCRE active in both the beta-1 and beta-2 subtype. This variant has significant heterogeneity in allelic imbalance in beta-2 and beta-2 chromatin accessibility, where the T2D risk allele has larger effect in beta-2 compared to beta-1. The values for allelic imbalance represent the fraction of reads from the risk allele and the 95% confidence interval. On the left is a schematic describing allelic imbalance mapping in reads from the beta-1 and beta-2 subtype. * P<.05. **(e)** Schematic showing abundance and functional changes of beta cell subtypes during T2D progression. The TFs maintaining beta cell subtype identity as well as TFs mediating T2D-associated changes are shown.

Given enrichment of T2D risk in cCREs defining the beta-1 and beta-2 subtypes, we next determined whether specific TFs that maintain subtype identity mediate this risk. Of the six TFs that maintain beta-1 and beta-2 subtype identity, genes encoding four of the TFs (*HNF1A, HNF4A, NEUROD1, TCF4*) harbor mutations known to cause Maturity Onset Diabetes of the Young (MODY), a monogenic form of diabetes^40^, and three of these TFs (*HNF1A, HNF4A, TCF4*) additionally map to known T2D risk loci^39^. We next determined whether subtype-defining binding sites for these TFs were enriched for T2D risk variants. There was significant enrichment for cCREs defining beta-1 identity bound by HNF4A and HNF4G (logOR=1.32, *P*=8.1×10^-3^; logOR=1.32, *P*=8.0×10^-3^) as well as nominal enrichment for cCREs bound by HNF1A (logOR=1.07, *P*=.033; **Figure 6b**). Similarly, there was significant enrichment for cCREs defining beta-2 identity bound by TCF4, NEUROD1 and NFIA (logOR=1.86, *P*=1.6×10^-4^; logOR=1.81, *P*=3.8×10^-4^; logOR=1.97, *P*=5.9×10^-4^). There was no corresponding evidence for enrichment (*P*>.05) in subtype-defining cCREs not bound by these TFs (**Figure 6b**).

In total there were 43 fine-mapped T2D variants that overlapped a cCRE defining beta-1 or beta-2 identity, including high-probability variants at the *GLIS3, RASGRP1, ZFPM1, SLC12A8, FAIM2,* and *SIX2/3* loci (**Supplementary Table 12**). We determined whether the T2D risk alleles of variants in cCREs defining beta-1 or beta-2 identity were correlated with increased or decreased chromatin accessibility using allelic imbalance mapping (Methods and **Supplementary Table 12**). Among fine-mapped T2D variants in cCREs defining beta-1 identity, T2D risk alleles were significantly more likely to reduce beta-1 accessibility than expected (obs=.86, exp.=.50, binomial *P*=0.013). We observed the same pattern among T2D-associated variants genome-wide in cCREs defining beta-1 identity (obs=.59, exp.=.50, binomial *P*=0.043). For example, at the 12p24 locus, rs1617434 overlapped a cCRE defining beta-1 identity where the T2D risk allele significantly (FDR<.10) decreased beta-1 accessibility (beta-1 allelic effect [π]=.27, 95% CI=.12,.46; q-value=.048) and was predicted to disrupt a HNF4A motif (**Figure 6c**). Furthermore, the same allele was associated with reduced expression of *ABCB9* (*P=*1.46×10^-7^), *RILPL2* (*P=*1.23×10^-6^), and *MPHOSPH9* (*P*=1.87×10^-3^), as well as other genes in islet expression QTL data^41^. By comparison, T2D risk alleles of variants in cCREs defining beta-2 identity were more likely to increase beta-2 accessibility than expected by chance (fine-mapped variants; obs=.67, exp.=.50, binomial *P*=.51; genome-wide variants; obs=.69, exp.=.50, binomial *P*=.011).

We finally identified T2D variants with heterogeneity in allelic effects on beta cell subtype activity that may modulate subtype identity. In total, we identified 163 fine-mapped T2D risk variants with at least nominal evidence for heterogeneity (*P*<.05) in beta-1 and beta-2 subtype chromatin accessibility (**Supplementary Table 13**). For example, at the 4q31 locus, fine-mapped T2D variant rs6813195 had heterogeneous effects on beta cell subtype chromatin accessibility (beta-1 π=.56, beta-2 π=.64, *P*=.024), where the T2D risk allele had increased accessibility in beta-2 compared to beta-1 cells (**Figure 6d**). The risk allele was also predicted to create a binding site for PAX6 and was associated with increased islet expression of *FBXW7* (*P*=7.49×10^-4^). In another example, at the 14q32 locus, fine-mapped T2D variant rs56330734 had heterogeneity in effects on beta cell subtype chromatin (beta-1 π=0, beta-2 π=.91, *P*=5.2×10^-5^). The T2D risk allele had increased accessibility in beta-2 compared to beta-1 cells and was predicted to create a NKX2-2 motif. In each of these examples, both the TFs and target genes affected by variant activity were involved in the NEUROD1-related GRN, suggesting that the variants may affect T2D risk by promoting beta-2 subtype identity.

Together, our analysis identifies two functionally distinct beta cell subtypes in human islets that are maintained by HNF1A, HNF4A and HNF4G and NEUROD1, TCF4 and NFIA, respectively (**Figure 6e**). We provide genetic evidence that the transcriptional programs maintaining beta cell subtype identity likely play a causal role in the pathogenesis of T2D. In T2D, there is an abundance shift between the two beta cell subtypes. Both subtypes are functionally impaired in T2D, and these functional changes are driven by signal-dependent TFs implicated in the cellular stress response.

## Discussion

Despite substantial efforts to define the molecular events underlying T2D pathogenesis in pancreatic islets, we still lack a thorough understanding of the gene regulatory programs driving T2D progression in beta cells and other islet cell types. Our study demonstrates the power of combining single-cell multiome data from a large sample number at different stages of disease with machine learning approaches, genetic association data, and single-cell functional measurements to define islet cell type and subtype gene regulatory programs involved in T2D pathogenesis. With the application of additional computational tools, our data can be further leveraged to improve fine-mapping of T2D risk loci, explore gene regulatory networks, and infer cell-cell interactions.

We used machine learning to identify beta cell subtypes and detected two beta cell subtypes in healthy donors which are functionally distinct and undergo a substantial abundance shift in T2D. Several studies have described beta cell subtypes based on cell surface markers^7^, gene expression^5^, chromatin activity^42^ and function using Patch-seq^16^. Core beta cell subtype-defining molecular features identified in our study are shared with those described in prior studies, indicating robustness of these subtypes across different cohorts and data types. For example, of the 28 most significant genes differentially expressed between beta subtypes based on cell surface marker expression^7^, 11 are differentially expressed between the two beta cell subtypes and another 11 genes showed the same sign of change albeit below our significance threshold. The same study^7^ also reported different insulin secretory activity of beta cell subtypes and an abundance shift in T2D concordant with our findings. A consistent observation across studies is the association of high insulin secretory capacity with high expression of insulin itself and genes involved in stimulus secretion coupling (e.g., *GCK, SYT1*). Our study expands prior studies by defining the GRNs that maintain the different beta cell subtypes. We show that feedback loops between TFs establish beta cell subtype identity. Specifically, we identify HNF1A, HNF4A and HNF4G as the core TFs maintaining the majority subtype in ND donors, whereas TCF4, NEUROD1 and NFIA maintain the minority subtype. Identification of these beta cell subtype-defining TFs can inform strategies for manipulating beta cell states for therapeutic intervention in T2D.

Our identification of two beta cell subtypes and their molecular and functional characterization in ND and T2D states provides novel insight into understanding T2D pathogenesis. Previous measurements of single-cell gene expression and exocytosis in beta cells by Patch-seq have shown that genes positively correlated with exocytosis in beta cells from ND donors are upregulated in T2D despite decreased exocytosis in T2D^16^. By identifying two distinct gene regulatory changes in T2D, our analysis provides a mechanistic understanding of this unexplained phenomenon. The most prominent gene regulatory change in T2D is an increase in the abundance of the beta cell subtype that in ND donors is the more highly exocytotic of the two subtypes, explaining why genes positively correlated with exocytosis are highly expressed in beta cells from T2D donors^16^. The second T2D-induced gene regulatory change occurs across both beta cell subtypes and is associated with decreased exocytosis. Thus, resolving beta cell subtypes allowed us to distinguish changes caused by the subtype shift from changes that occur in all beta cells in T2D.

The T2D-induced gene expression changes across both beta cell subtypes are driven by signal-dependent TFs, many of which (e.g., XBP1, ATF6, TFEB, and DBP) are downstream effectors of the ER stress and integrated stress response^30–32^. Experimental models provide evidence that these TFs are regulated by elevated glucose and free fatty acid levels^33^, indicating that the activity change of these TFs in beta cells from T2D donors is likely a consequence of T2D-associated metabolic abnormalities. Downstream of these TFs, we identify a network of genes involved in processes relevant for beta cell function, including protein translation and protein quality control, oxidative phosphorylation, and vesicle trafficking. Given evidence from *in vitro* models that high glucose and free fatty acids impair beta cell function and lead to similar gene expression changes^33^ as we observed in T2D, the identified “stress response GRN” likely causes impaired exocytosis in T2D. This view is further supported by evidence that decreased DBP^31^ and XBP1^43^ or increased ATF6^44^ activity impair beta cell function. Our findings suggest that reversal of the changes induced by these signal-dependent TFs will be essential for reversing beta cell dysfunction in T2D.

The most prominent change in T2D is the shift from the beta cell subtype that is less exocytotic to the one that is more exocytotic in ND individuals. This raises the question of whether the subtype shift represents a compensatory mechanism early in disease or whether it contributes to T2D pathogenesis. Several observations support the view that the subtype shift could have a causal rather than compensatory role in disease. First, our genetic evidence supports causality for T2D. We found that active chromatin distinguishing the two beta cell subtypes is preferentially enriched for T2D risk variants compared to general beta cell active chromatin. In addition, among T2D variants in active chromatin defining the less exocytotic subtype, the T2D risk alleles are correlated with reduced chromatin activity, indicating that T2D risk alleles favor a transition toward the more exocytotic beta cell subtype. Further arguing against a compensatory role of the beta cell subtype shift, we observed no significant difference in beta cell subtype composition between pre-T2D and ND donors, suggesting that the shift is a later event in disease progression and not present early when compensatory mechanisms might operate.

We identify HNF4A as central to the GRN that defines the less secretory beta cell subtype and show enrichment of T2D risk variants in HNF4A binding sites in this subtype, suggesting that reduced HNF4A activity could trigger a shift toward the more exocytotic beta cell subtype. *HNF4A* loss-of-function mutations cause MODY-1 in humans, which is characterized by early insulin hypersecretion followed by progression to beta cell dysfunction and diabetes later in life^40^. Thus, our results and clinical findings in MODY-1 patients support a mechanism whereby loss of HNF4A activity could be a causal event in T2D pathogenesis leading to increased insulin secretion. How a shift toward a more secretory beta cell subtype leads to beta cell failure is still an open question. It is possible that the beta cell subtype-defining GRN and the T2D-induced “stress response GRN” are intricately linked and that both gene regulatory changes occur simultaneously during T2D progression. This view is supported by evidence showing that loss of HNF1A function reduces XBP1 and sensitizes beta cells to ER stress^45^. Conversely, genetic reduction of insulin dosage - akin of forcing beta cells into a less exocytotic subtype - alleviates beta cell ER stress^46^. Therefore, the more highly exocytotic beta cell subtype may ultimately be more vulnerable and prone to fail in the face of metabolic stress. However, given the heterogeneity of human islet samples, it will be important to validate inferences made from the GRNs on additional human islet data sets.

Another major advance of our study is the development of a classifier based on machine learning to identify disease-associated patterns in single-cell data. The heterogeneity of human samples imposes challenges for analyzing and interpreting single-cell data from primary human tissues. We demonstrate that our classifier robustly identifies cell subtypes across different human islet data sets. Notably, these subtypes could not be identified by standard and widely used unsupervised dimensionality reduction methods likely due to donor-specific confounding factors. The machine learning approach presented here should have broad applications for identifying disease-relevant patterns in single-cell data also from other primary human tissues.

## Methods

### Human islets

We obtained islet preparations for 34 donors from 4 resource centers (22 from City of Hope National Medical Center, 9 from Scharp-Lacy Research Institute, 2 from the University of Pennsylvania, and 1 from the University of Wisconsin). Characteristics (i.e., age, sex, BMI, HbA1c, ethnicity) and available clinical information for individual donors are listed in Supplementary Table 1a. The mean age, BMI, and HbA1c, as well as number of donors by sex and ethnicity in each disease group are summarized in Supplementary Table 1b. Classification of donors as non-diabetic (ND), pre-T2D or T2D was based on the person’s medical record or post-mortem HbA1c value. Donors with prior T2D diagnosis per medical record or HbA1c ≥ 6.5 were classified as T2D, donors without prior T2D diagnosis and 5.7 ≤ HbA1c ≤ 6.4 as pre-T2D, and donors without prior T2D diagnosis and HbA1c ≤ 5.6 (or HbA1c unavailable) as ND. Islet preparations were further enriched using zinc-dithizone staining followed by hand picking, and snap frozen with liquid nitrogen or dry ice. Studies were given exempt status by the Institutional Review Board (IRB) of the University of California San Diego.

### Generation of snATAC-seq data using the 10x Chromium platform

Approximately 1,000 islet equivalents (∼1,000 cells per IEQ) were resuspended in 1 mL nuclei permeabilization buffer (10 mM Tris-HCL (pH 7.5), 10 mM NaCl, 3mM MgCl2, 0.1% Tween-20 (Sigma), 0.1% IGEPAL-CA630 (Sigma), 0.01% Digitonin (Promega) and 1% fatty acid-free BSA (Proliant 68700) in molecular biology-grade water) and homogenized using 1 mL glass dounce homogenizer with a tight-fitting pestle (Wheaton, EF24835AA) for 10-20 strokes until the solution was homogeneous. Homogenized islets were filtered with 30 µm filter (CellTrics, Sysmex) and then incubated for 10 min at 4°C on a rotator. Nuclei were pelleted with a swinging bucket centrifuge (500 x g, 5 min, 4°C; 5920R, Eppendorf) and washed with Wash buffer (10 mM Tris-HCL (pH 7.5), 10 mM NaCl, 3 mM MgCl_2_, 0.1% Tween-20, and 1% BSA (Proliant 68700) in molecular biology-grade water). Nuclei were pelleted and resuspended in 30 µL of 1x Nuclei Buffer (10x Genomics). Nuclei were counted using a hemocytometer, and 15,360 nuclei were used for tagmentation. Single-cell ATAC-seq libraries were generated using the Chromium Single Cell ATAC Library & Gel Bead Kit (10x Genomics, 1000110), Chromium Chip E Single Cell ATAC kit (10x Genomics, 1000086) and indexes (Chromium i7 Multiplex Kit N, Set A, 10x Genomics, 1000084) following manufacturer instructions. Final libraries were quantified using a Qubit fluorimeter (Life Technologies) and the nucleosomal pattern was verified using a Tapestation (High Sensitivity D1000, Agilent). Libraries were sequenced on NextSeq 500, HiSeq 4000 and NovaSeq 6000 sequencers (Illumina) with following read lengths: 50 + 8 + 16 + 50 (Read1 + Index1 + Index2 + Read2).

### Generation of joint single nucleus RNA and ATAC-seq data using Chromium Single-cell Multiome ATAC + Gene Expression (10x Genomics)

Islets were resuspended in 1 mL wash buffer (10mM Tris-HCL (pH 7.4), 10mM NaCl, 3mM MgCl2, 0.1% Tween-20 (Sigma), 1% fatty acid-free BSA (Proliant, 68700), 1 mM DTT (Sigma), 1x protease inhibitors (Thermo Fisher Scientific, PIA32965), 1U/µl RNAsin (Promega, N2515) in molecular biology-grade water) and homogenized using 1 mL glass dounce homogenizer with a tight-fitting pestle (Wheaton, EF24835AA) for 10-20 strokes until the solution was homogeneous. Homogenized islets were filtered with 30 µm filter (CellTrics, Sysmex) and pelleted with a swinging bucket centrifuge (500 x g, 5 min, 4°C; 5920R, Eppendorf). Nuclei were resuspended in 400 µL of sort buffer (1% fatty acid-free BSA, 1x protease inhibitors (Thermo Fisher Scientific, PIA32965), 1U/µl RNAsin (Promega, N2515) in PBS) and stained with 7-AAD (1 µM; Thermo Fisher Scientific, A1310). 120,000 nuclei were sorted using an SH800 sorter (Sony) into 87.5 μl of collection buffer (1U/µl RNAsin (Promega, N2515), 5% fatty acid-free BSA (Proliant, 68700) in PBS). Nuclei suspension was mixed in a ratio of 4:1 with 5x permeabilization buffer (50 mM Tris-HCL (pH 7.4), 50 mM NaCl, 15 mM MgCl2, 0.5% Tween-20 (Sigma), 0.5% IGEPAL-CA630 (Sigma), 0.05% Digitonin (Promega), 5% fatty acid-free BSA (Proliant, 68700), 5 mM DTT (Sigma), 5x protease inhibitors (Thermo Fisher Scientific, PIA32965), 1U/µl RNAsin (Promega, N2515) in molecular biology-grade water) and incubated on ice for 1 min before pelleting with a swinging-bucket centrifuge (500 x g, 5 min, 4°C; 5920R, Eppendorf). Supernatant was gently removed and ∼50 µl were left behind to increase nuclei recovery. 650 µl of wash buffer (10mM Tris-HCL (pH 7.4), 10mM NaCl, 3mM MgCl2, 0.1% Tween-20 (Sigma), 1% fatty acid-free BSA (Proliant, 68700), 1 mM DTT (Sigma), 1x protease inhibitors (Thermo Fisher Scientific, PIA32965), 1U/µl RNAsin (Promega, N2515) in molecular biology-grade water) were added without disturbing the pellet and nuclei were pelleted with a swinging bucket centrifuge (500 x g, 5 min, 4°C; 5920R, Eppendorf). Supernatant was gently removed without disturbing the pellet and leaving ∼2-3 µl behind. 7-10 µl of 1x Nuclei Buffer (10x Genomics) was added and nuclei gently resuspended. Nuclei were counted using a hemocytometer, and 16,550-18,000 nuclei were used as input for tagmentation. Single-cell Multiome ATAC + Gene Expression libraries were generated following manufacturer instructions (Chromium Next GEM Single-cell Multiome ATAC + Gene Expression Reagent Bundle, 1000283; Chromium Next GEM Chip J Single cell, 1000234; Dual Index Kit TT Set A, 1000215; Single Index Kit N Set A, 1000212; 10x Genomics) with these PCR cycles: 7 cycles for ATAC index PCR, 7 cycles for cDNA amplification, 13-16 cycles for RNA index PCR. Final libraries were quantified using a Qubit fluorimeter (Life Technologies) and the size distribution was checked using a Tapestation (High Sensitivity D1000, Agilent). Libraries were sequenced on NextSeq 500 and NovaSeq 6000 sequencers (Illumina) with following read lengths (Read1 + Index1 + Index2 + Read2): ATAC (NovaSeq 6000) 50 + 8 + 24 + 50; ATAC (NextSeq 500 with custom recipe) 50 + 8 + 16 + 50; RNA (NextSeq 500, NovaSeq 6000): 28 + 10 + 10 + 90.

### Raw data processing and quality control

#### Data processing using Cell Ranger ATAC and ARC software

Alignment to the hg19 genome and initial processing were performed using the 10x Genomics Cell Ranger ATAC v1.1.0 and multiome ARC v.2.0.0 pipelines. We filtered reads with MAPQ<30, secondary or unmapped reads, and duplicate reads from the resulting bam files using samtools^47^. Sample information and a summary of the Cell Ranger ATAC-seq and mutiome quality metrics are provided in **Supplementary Table 1a**.

#### Filtering barcode doublets and low-quality cells for each individual donor

Cell barcodes from the 10x Chromium snATAC-seq assay may have barcode multiplets that have more than one oligonucleotide sequence^48^. We used ‘clean_barcode_multiplets_1.1.py’ script from 10x to identify barcode multiplets for each donor and excluded these barcodes from further analysis. We then filtered low quality snATAC-seq profiles by total UMIs (<1,000), fraction of reads overlapping TSS (<15%), fraction of reads overlapping called peaks (<30%), and fraction of reads overlapping mitochondrial DNA (>10%) according to the distribution of these metrics for all barcodes. We also excluded profiles that had extremely high unique nuclear reads (top 1%), fraction of reads overlapping TSS (top 1%) and called peaks (top 1%) to minimize the contribution of these barcodes to our analysis. Representative cell filtering from donor JYH809 is shown in **Supplementary Figure 1b**. For multiome data, we used identical cutoffs to filter cells with low quality ATAC profiles and used total UMIs (<1,000) and fraction of reads overlapping mitochondrial DNA (>10%) to filter cells with low quality RNA profiles.

#### Cell clustering

After filtering low quality cells, we checked data quality from each sample by performing an initial clustering using Scanpy (v.1.6.0)^49^. We partitioned the hg19 genome into 5 kb sliding windows and removing windows overlapping blacklisted regions from ENCODE^50, 51^ (https://www.encodeproject.org/annotations/ENCSR636HFF/). Using 5 kb sliding windows as features, we produced a barcode-by-feature count matrix consisting of the counts of reads within each feature region for each barcode. We normalized each barcode to a uniform read depth and extracted highly variable windows. Then, we regressed out the total read depth for each cell, performed PCA, and extracted the top 50 principal components to calculate the nearest 30 neighbors using the cosine metric, which were subsequently used for UMAP dimensionality reduction with the parameters ‘min_dist=0.3’ and Leiden^52^ clustering with the parameters ‘resolution=0.8’. Representative cell clustering and marker gene promoter accessibility from donor JYH809 are shown in **Supplementary Figure 1c,d**.

We then performed initial cell clustering for 255,598 cells from all donors using similar methods to cluster cells for each donor. Of note, we extracted highly variable windows across cells from all experiments. Since read depth was a technical covariate specific to each experiment, we regressed this out on a per-experiment basis. We also used Harmony^53^ to adjust for batch effects across experiments.

We identified clusters and subclusters (‘resolution’=1.5) with significantly different total UMIs, fraction of reads overlapping TSS, or fraction of reads overlapping called peaks compared to other clusters and subclusters. We excluded these clusters and subclusters from further analysis, exemplified in by cluster 14 and subcluster 1 from cluster 6 in **Supplementary Figure 1f**. We also used marker hormones for alpha (*GCG*), beta (*INS-IGF2*), and delta (*SST*) cells to identify and remove potential doublets that have chromatin accessibility in more than one marker gene promoter. We retained 218,973 barcodes after excluding 22,929 cells in low-quality clusters and subclusters (8.9%) and 13,696 potential doublets (5.3%) and used identical methods to cluster these retained barcodes. UMAPs for cell clustering and marker gene promoter accessibility are shown in **Supplementary Figure 1g,h**.

We aggregated reads within each cluster (**Supplementary Figure 1e**) and called peaks for each cluster using the MACS2 call peak command with parameters ‘--nomodel --extsize 200 –shift 0 --keep-dup all -q 0.05’ and filtered these peaks by the ENCODE hg19 blacklist. Then, we merged peaks from all clusters to get a union peak set containing the peaks observed across all clusters. We used these union peaks as features to generate a barcode-by-feature count matrix consisting of the counts of reads within each feature region for each barcode. We performed cell clustering using identical methods for initial clustering of all cells and identified 13 cell clusters (Figure 1b). We determined the cell type represented by each cluster by examining chromatin accessibility at the promoter regions of known marker genes for alpha (*GCG*), beta (*INS-IGF2*), delta (*SST*), gamma (*PPY*), acinar (*REG1A*), ductal (*CFTR*), stellate (*PDGFRB*), endothelial (*CLEC14A*), and immune cells (*CCL3*).

### Generating fixed-width and nonoverlapping peaks that represent open chromatin sites across all cell types

We called peaks for each cell type in Figure 1b using the MACS2 call peak command with parameters ‘--nomodel --extsize 200 –shift 0 --keep-dup all -q 0.05’ and filtered these peaks by the ENCODE hg19 blacklist. For each cell type, we generated fixed-width peaks (summits of these peaks from macs2 were extended by 250 bp on either side to a final width of 501 bp), as previously described^54^. We quantified the significance of these fixed-width peaks in each cell type by converting the MACS2 peak scores (−log10(Q value)) to a ‘score quantile’. Then, fixed-width peaks for each cell type were combined into a cumulative peak set. As there are overlapping peaks across cell types, we retained the most significant peak and any peak that directly overlapped with that significant peak was removed. This process was iterated to the next most significant peak and so on until all peaks were either kept or removed due to direct overlap with a more significant peak. In total, we got 412,113 fixed-width (501 bp) and nonoverlapping peaks. By identifying fixed-width peaks that have overlap with peaks for each cell type from MACS2, we got fixed-width peaks for alpha (246,919 peaks), beta (230,573 peaks), delta (168,925 peaks), gamma (121,170 peaks), acinar (157,284 peaks), ductal (135,264 peaks), EC (81,953 peaks), immune (87,203 peaks), and stellate cells (120,114 peaks).

### Identification of beta cell subtypes using machine learning

#### Train and test classifier to distinguish beta cells from different disease states

We used chromatin accessibility of 224,563 beta cell autosomal cCREs to characterize individual beta cells. 90,290 beta cells (35,103 beta cells from 11 ND, 19,682 beta cells from pre-T2D, 35,505 beta cells from T2D donors) were retained after excluding beta cells with less than 1,000 reads within beta cell autosomal cCREs. We used beta cells from one donor at a time as a testing group while using beta cells from remaining donors as a training group (**Supplementary Figure 5c**). Using the chromatin accessibility profiles of training beta cells and their disease state annotation, we trained a classifier using XGBOOST^20^ (v.0.80.1) to distinguish beta cells from ND, pre-T2D and T2D donors. We then predicted the disease state of beta cells from donors in the testing group using the trained classifier and compared predictions to the annotated disease state of testing donors to calculate the prediction accuracy. We used each donor as a testing group and obtained prediction accuracies for each donor. We down-sampled beta cells from ND and T2D donors to numbers from pre-T2D donors and repeated the training and testing steps to test the effect of cell numbers.

### Train classifier to predict two beta cell subtypes

After recognizing two major beta cell subtypes enriched in either ND (beta-1 subtype) or T2D (beta-2 subtype) donors, we used reiterative training and testing steps to obtain a classifier distinguishing the two beta cell subtypes (**Supplementary Figure 5j**). Using beta cells from ND (11 donors, 35,103 beta cells) and T2D (15 donors, 35,505 beta cells) donors, we trained and tested the classifier as described above. Since beta-1 and beta-2 cells coexisted in each donor, we used reiterative model training and testing to identify the dominant beta cell subtype in ND (beta-1) and T2D (beta-2) donors. For each round of training and testing, we used beta cells whose disease state was correctly predicted for the next round of training and testing until the disease state of all selected beta cells was correctly predicted. Using this methodology, we obtained the final classifier to distinguish beta-1 and beta-2 cells and used the classifier to predict subtype identity of beta cells from pre-T2D donors in our snATAC-seq data and in an independent islet snATAC-seq dataset from ND and T2D donors from the Human Pancreas Analysis Program (HPAP) (see below).

### Computing co-accessibility using Cicero

For each endocrine cell type, we used Cicero^55^ (v.1.3.4.10) to calculate co-accessibility scores for pairs of peaks for alpha, beta, delta, and gamma cells. We started from the merged peak by cell sparse binary matrix, extracted alpha cells, and filtered out peaks that were not present in alpha cells. We used the ‘make_cicero_cds’ function to aggregate cells based on the 50 nearest neighbors. We then used Cicero to calculate co-accessibility scores using a window size of 1 Mb and a distance constraint of 250 kb. We then repeated the same procedure for beta, delta, and gamma cells. We used a co-accessibility threshold of 0.05 to define pairs of peaks as co-accessible. Peaks within and outside ± 5 kb of a TSS in GENCODE V19 were considered proximal and distal, respectively. Peaks within ± 500 bp of a TSS in GENCODE V19 were defined as promoter. Co-accessible pairs were assigned to one of three groups: distal-to-distal, distal-to-proximal and proximal-to-proximal. Distal-to-proximal co-accessible pairs were defined as potential enhancer-promoter connections. Genes linked to proximal or distal cCREs were identified.

### Differential peak and gene expression analysis

#### Identification of independent confounding factors in snATAC-seq data using PCA

To determine the factors that account for sample variability in our data, we conducted principle component analysis (PCA) on cell type-specific pseudo-bulk profiles generated from each of the 34 donors. Here, features were fixed-width peaks for each cell type and donor. Next, we calculated total-count normalized matrices, applied PCA to the normalized matrices using prcomp in R, and visualized the position of each donor using the autoplot function in R. In addition to disease status (ND, pre-T2D, T2D), we considered HbA1c, age, body mass index (BMI), and sex as biological covariates as well as islet index, islet purity, sequencing depth, total read counts, and the fraction of reads overlapping TSS as technical covariates. We calculated the absolute Spearman correlation coefficient between the first 6 PCs and each biological or technical variable. We used an absolute Spearman correlation threshold of 0.4 as a cutoff to identify factors that have high correlation with each PC. We further identified independent confounding factors by calculating the pairwise Spearman correlation coefficients between factors. As high pairwise association (Spearman’s ρ >0.9) represents dependencies between factors such as disease status and HAb1c level, we only retained one of them. In beta cells, we found a high correlation of the fraction of reads overlapping TSS with PC1; the islet index with PC2; disease status, Hba1c, and total read counts with PC3, disease status and Hba1c with PC4; and the fraction of reads overlapping TSS with PC5 (**Supplementary Figure 2a,b**). Calculation of the pairwise Spearman correlation coefficients between variates revealed a high degree of correlation between interdependent variables, such as HAb1c levels and disease status, and identified the fraction of reads overlapping TSS, the islet index, and total read counts as independent confounding factors in our data (**Supplementary Figure 2c**). We obtained similar results for alpha, delta, and gamma cells (**Supplementary Figure 2d-l**).

#### Identification of differential peaks in cell type pseudo-bulk data with DESeq2

For each cell type, we called differential peaks between disease groups (i.e., pre-T2D vs ND, T2D vs pre-T2D and T2D vs ND) using DESeq2^19^ in the R package. We used the cell type-specific pseudo-bulk feature-by-donor matrix (11 ND, 8 pre-T2D and 15 T2D donors) as input and major biological and technical confounding factors (age, BMI, sex, islet index, fraction of reads overlapping TSS, and total reads) as covariates. An FDR <0.1 (p-values adjustment with the Benjamini-Hochberg method) was used as the cutoff to identify differential peaks. We also identified differential peaks based on age, sex, and BMI. We used CEAS^56^ to annotate differential sites. Of note, we found very few (0-301) differential peaks in each islet cell type based on sex, age, and BMI, suggesting no consistent effect on chromatin accessibility in our data. We performed down-sampling to match cell numbers for alpha, beta and delta cells. We down-sampled alpha, beta, delta cells by randomly selecting 15,000 and 5,000 cells. Then, we called differential cCREs using down-sampled cells. We also performed down-sampling to match donor numbers in the ND, pre-T2D and T2D groups. We down-sampled ND and T2D donors by randomly selecting 8 donors from all ND and T2D donors. Then, we called beta cell differential cCREs with identical sample size (n=8) for ND, pre-T2D and T2D groups. We repeated this process by randomly selecting six different combinations of 8 ND and T2D donors.

#### Identification of differential peaks and genes between beta cell subtypes using paired t-test

We generated beta-1 and beta-2 pseudo-bulk accessibility profiles (34 total, *n* = 11 ND, *n* = 8 pre-T2D, *n* = 15 T2D donors) from snATAC-seq data and gene expression profiles from multiome data (20 total, *n* = 6 ND, *n* = 8 pre-T2D, *n* = 6 T2D donors). Using these pseudo-bulk profiles, we performed paired t-test to identify differential cCREs (FDR<.05, p-values adjusted with the Benjamini-Hochberg method) and genes (FDR<.15, p-values adjusted with the Benjamini-Hochberg method) between beta cell subtypes. We calculated the Pearson correction between log_2_ differences (beta-2/beta-1) in chromatin accessibility at differential cCREs and log_2_ differences (beta-2/beta-1) in gene expression of cCRE target genes with differential expression. To identify high confidence differentially expressed genes between beta cell subtypes, we only focused on differential expressed genes that also have significant changes in proximal (within ± 5 kb of a TSS in GENCODE V19) or distal cCREs accessibility (defined in “Computing co-accessibility and identifying distal cCREs using Cicero” section) between beta cell subtypes.

### TF motif enrichment analysis

Using the barcode-by-peaks (501 bp fixed-width peaks) count matrix as input, we inferred enrichment of TF motifs for each barcode using chromVAR^57^ (v.1.4.1). We filtered cells with minimal reads less than 1500 (min_depth=1500) and peaks with fraction of reads less than 0.15 (min_in_peaks=0.15) by using ‘filterSamplesPlot’ function from chromVAR. We also corrected GC bias based on ‘BSgenome.Hsapiens.UCSC.hg19’ using the ‘addGCBias’ function. Then, we used the TF binding profiles database JASPAR 2020 motifs^58^ and calculated the deviation z-scores for each TF motif in each cell by using the ‘computeDeviations’ function. High-variance TF motifs across all cell types were selected using the ‘computeVariability’ function with the cut-off 1.15 (*n*=255). For each of these variable motifs, we calculated the mean z-score for each cell type and normalized the values to 0 (minimal) and 1 (maximal).

We performed both *de novo* and known motif enrichment analysis using HOMER^59^ (v.4.11) command ‘findMotifsGenome.pl’. We focused on significantly enriched *de novo* motifs and assigned the best matched known TF motifs to *de novo* motifs.

### Gene ontology enrichment analysis

We performed gene ontology enrichment analysis using R package Enrichr^60^. Library “GO_Biological_Process_2018” was used with default parameters.

### Inferring gene regulatory networks from multiome data

We first used a position frequency matrix (PFMatrixList object) of TF DNA-binding preferences from the JASPAR 2020 database^58^ and width-fixed peaks as input to perform TF binding motif analysis. We used the ‘matchMotifs’ function in the R package motifmatchr to infer beta cell cCREs occupied by 264 TFs expressed in beta cells (mean TPM across donors >4). We linked beta cell cCREs occupied by each TF to target genes based on proximity to the gene promoter (within ± 5 kb of a TSS in GENCODE V19) or co-accessibility between the distal cCRE and gene promoter across single beta cells (defined in “Computing co-accessibility and identifying distal cCREs using Cicero” section). We further calculated gene expression correlations between each TF and its target genes in aggregate beta-1 and beta-2 cells for each donor from multiome data (*n*=20 donors). For each TF, we identified target genes that have significant positive and negative gene expression Pearson correlation with the TF (FDR<0.05, p-values adjusted with the Benjamini-Hochberg method) and defined positively correlated TF-gene modules and negatively correlated TF-gene modules.

### Identification of differential TF-gene modules

We performed gene set analysis using R package GSA^23^ (v.1.3.1) to evaluate changes of individual TF-gene modules (using all genes in the TF-gene module) between beta cell subtypes and during T2D progression (20 total, *n*=6 ND, *n*=8 pre-T2D, *n*=6 T2D donors, each donor has beta-1 and beta-2 pseudo-bulk gene expression profiles). We used a p-value<0.05 and enrichment score to identify significantly up (enrichment score>0.6) or down (enrichment score < −0.6) regulated TF-gene modules between beta cell subtypes. We further filtered these TF-gene modules by intersecting with enriched TF motifs in cCREs with higher accessibility in beta-1 or beta-2. For each beta cell subtype, we used a p-value<0.05 and enrichment score to identify significantly up (enrichment score>1.3) or down (enrichment score < −1.3) regulated TF-gene modules during T2D progression. We further filtered the TFs by intersecting with enriched TF motifs in cCREs with significant changes in beta-1 or beta-2 during T2D progression.

### Public human islet snATAC-seq and scRNA-seq data

We downloaded public human islet snATAC-seq data from Human Pancreas Analysis Program (HPAP, https://hpap.pmacs.upenn.edu/; V2.0.0, data download date: 07/09/2021). We processed and analyzed the data using the pipeline described above. After quality control, snATAC-seq data were used to validate results from our snATAC-seq data. Donor characteristics are summarized in Supplementary Table 14a. More information about these donors is available via https://hpap.pmacs.upenn.edu/explore/donor?by_donor.

We downloaded scRNA-seq data and metadata of donors from three public islet scRNA-seq datasets^5, 12, 22^. We processed and analyzed the data using the pipeline described above. Donor characteristics are available in the original publications and summarized in Supplementary Table 14b-d.

To classify donors from public islet datasets analyzed in this study as ND, pre-T2D or T2D we applied the same classification criteria as used for classifying the 34 donors from the cohort profiled in this study (see “Human islets”). In some cases, our classification criteria differed from the criteria used in the original studies leading to reclassification of select donors (see Supplementary Table 14).

### GWAS enrichment analysis

We tested for enrichment of fine-mapped T2D risk variants from the DIAMANTE consortium for beta cell cCREs defining the beta-1 and beta-2 subtype as well as cCREs with differential activity in T2D. For each set of cCREs, we calculated the cumulative posterior probability of association (cPPA) of all fine-mapped variants overlapping cCREs. We then generated a null distribution of cPPA by randomly selecting the same number of cCREs from the set of all beta cell cCREs across 100,000 permutations. We calculated a p-value as the number of permutations with a higher cPPA than for the observed set of cCREs. We further computed an odds ratio as cPPA_obs_*(cPPA_max_-cPPA_mean_)/cPPA_mean_*(cPPA_max_-cPPA_obs_), where cPPA_obs_ was the observed cPPA, cPPA_max_ is the maximum possible cPPA for that number of sites and cPPA_mean_ is the average cPPA from the null distribution, and took the natural log of the odds ratio.

### Genotyping and imputation

1000-3000 IEQ human islets pellets were resuspended in 200 µL PBS and treated with 20 µL 10 mg/mL Rnase A (Invitrogen) and 20 µL Protein Kinase K (Qiagen) for 30 min at RT followed by the steps as described in the protocol of Dneasy Blood & Tissue Kit (QIAGEN). 200-500 ng DNA was used for genotyping using the Infinium Omni2.5-8v1-4 and the Infinium Omni2.5-8v1-5 Genotyping BeadChip (Illumina) at the UCSD IGM core. We called genotypes with GenomeStudio (v.2.0.4) using default settings. For genotypes that passed quality filters (missing<0.05, minor allele frequency (MAF>0.01), non-ambiguous alleles defined by AT/GC variants with MAF>40%), we imputed genotypes into the TOPMed r2 reference panel^61^ using the TOPMed Imputation Server^62^. Post-imputation, we removed genotypes with low imputation quality (R^2^<0.3) and used liftOver^63^ to map the coordinates back to hg19.

### Allelic imbalance analysis

To estimate cell type-specific chromatin accessibility allelic imbalance (AI), we modified the WASP^64^ pipeline for single-cell analysis by re-mapping reads using phase information and removing duplicate reads within each cell. For each sample, we aggregated re-mapped reads for cells from each beta cell subtype. We assessed AI at each heterozygous variant using a binomial test, assuming a null hypothesis of equal proportions of reads for each allele. We meta-analyzed z-scores across all samples using Stouffer’s z-score method with re-mapped read depth as a weight. We used AI z-scores to calculate 2-sided p-values. We annotated fine-mapped T2D variants in 99% credible sets from DIAMANTE^65^ overlapping cCREs defining beta cell subtype identity with AI z-scores, and calculated q-values for these variants using Storey’s method (R package qvalue v2.16.0). For each subtype, we identified the most probable fine-mapped variant per T2D signal overlapping cCREs defining identity of that subtype. We then determined whether the proportion of T2D risk alleles for these variants with decreased subtype accessibility differed from the expected proportion of .50 using a binomial test. We further identified all variants with *P*<.0001 in DIAMANTE^65^ overlapping cCREs defining beta cell subtype identity, and again determined whether the proportion of T2D risk alleles for these variants with decreased accessibility differed from the expected proportion using a binomial test.

For the analyses comparing AI between beta cell subtypes, we retained variants tested for AI in at least two samples for each subtype and used two-sided binomial proportion tests to compare AI z-scores between subtypes. We obtained islet eQTL data from the TIGER database (tiger.bsc.es).

### Data availability

Single nucleus ATAC sequencing data and processed data are available through the Gene Expression Omnibus under accession GSE169453, and single nucleus multiome data under accession GSE200044 and genotyping data under accession GSE170763. UCSC genome browser sessions of aggregated snATAC-seq data are available at: https://genome.ucsc.edu/s/gaowei/hg19_cell_type, https://genome.ucsc.edu/s/gaowei/hg19_beta_cell. Previously published^16, 17^ Patch-seq data are available as raw sequencing reads in NCBI GEO under accession numbers GSE124742 and GSE164875. Additional Patch-seq data are accessible at the HPAP database URL- https://hpap.pmacs.upenn.edu.

### Code availability

Custom codes for main analysis used in this study have been deposited on GitHub: https://github.com/gaoweiwang/Islet_snATACseq.

## Acknowledgements

This publication includes data generated at the UC San Diego IGM Genomics Center utilizing an Illumina NovaSeq 6000 that was purchased with funding from a National Institutes of Health SIG grant (#S10 OD026929). This work was supported by NIH U01DK105541 and R01DK122607 to M.S. and K.G., R01DK114650 to K.G., R01DK068471 to M.S, U01DK120447 to P.E.M., U01DK123716 to S.K.K. and P.E.M. and UC4-DK112217, UC4-DK112232, and P30 DK116074 to S.K.K. Work at the UCSD Center for Epigenomics was supported by the UC San Diego School of Medicine. We thank Hong Gao, Yi Shi, Kelly A. Frazer, Bing Ren, and members of Sander lab for scientific discussions and input on the project and the organ donors and their families for their contributions to make this study possible.

## Author Contributions

M.S., K.J.G. and S.P. conceived and supervised the research in the study; M.S., K.J.G., G.W., and J.C. wrote the manuscript; G.W. and J.C. performed analyses of single-cell and genetic data; C.Z., I.M. N.K., J.Y.H, and M.L.O. performed experiments; M.Mi. performed 10x single-cell assays; E.B. and M.Ma. contributed to data analyses. F.R.K. provided human islets. J.C-S., T.dS., XQ.D., C.E., Y.H., S.K.K., and P.E.M. provided Patch-seq data.

## Supplementary Figures

**Supplementary Figure 1.**
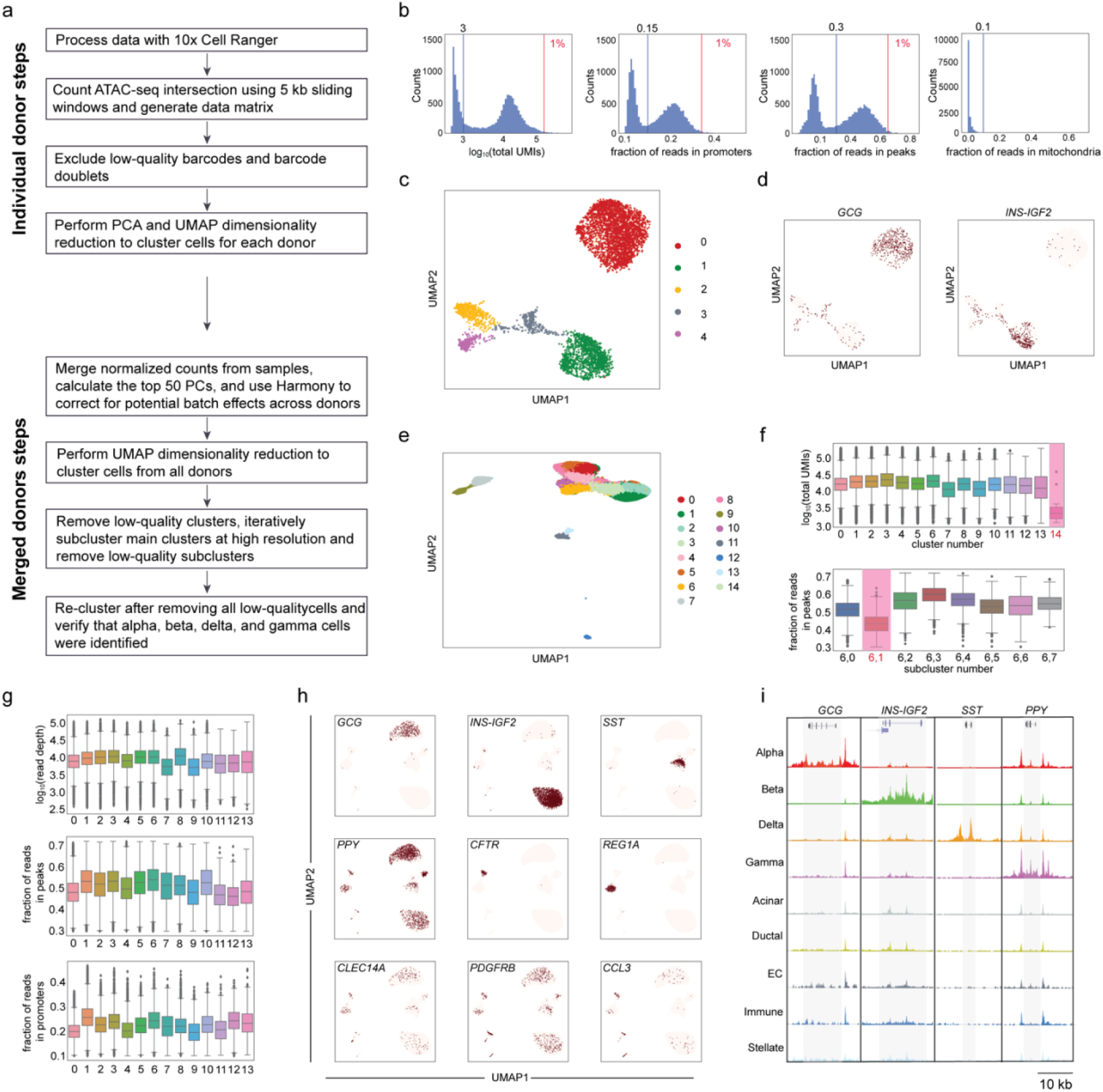
Quality control of snATAC-seq data. **(a)** Steps for snATAC-seq data processing and quality control. **(b)** Representative quality control (QC) metrics for each donor. Log_10_ total UMIs, fraction of reads overlapping promoters, fraction of reads overlapping peaks, and fraction of reads overlapping mitochondria DNA distribution of cells from T2D donor JYH809 as example. Blue vertical lines denote thresholds of 1000 minimal fragment number, 15% fragments overlapping promoters, 30% fragments overlapping peaks, and 10% fraction of reads overlapping mitochondria DNA, respectively. Red vertical lines denote thresholds to identify top 1% barcodes with extremely high total fragment number and fraction of reads overlapping promoters and peaks, respectively. **(c)** Representative cell clustering from donor JYH809 conducted for each donor. Cells are plotted using the first two UMAP components. **(d)** Promoter chromatin accessibility in a 5 kb window around TSS for endocrine marker genes for each profiled cell from donor JYH809. Total counts normalization and log-transformation were applied. **(e)** Cell clustering of chromatin accessibility profiles from all donors. Cells are plotted using the first two UMAP components. **(f)** Representative low-quality cluster and subcluster. Log_10_ total UMIs distribution of cells from each cluster. Cells in cluster 14 (top, highlighted in red) have significantly lower unique fragment than cells in other clusters. Fraction of reads overlapping peaks distribution of cells from each subcluster of main cluster 6. Cells in subcluster 1 (bottom, highlighted in red) have significantly lower fraction of reads overlapping peaks than cells in other clusters. **(g)** Log_10_ total UMIs, fraction of reads overlapping peaks and fraction of reads in promoters of cells from each cluster in Figure 1b, showing that these metrics do not drive single-cell grouping in UMAP space. **(h)** Promoter chromatin accessibility in a 5 kb window around TSS for selected endocrine and non-endocrine marker genes for each profiled cell (alpha: *GCG*, beta: *INS-IGF2*, delta: *SST*, gamma: *PPY*, acinar: *REG1A*, ductal: *CFTR*, stellate: *PDGFRB*, endothelial: *CLEC14A*, immune: *CCL3*). Total counts normalization and log-transformation were applied. **(i)** Genome browser tracks showing aggregate read density (scaled to uniform 1×10^6^ read depth) for cells within each cell type cluster at hormone gene loci for endocrine islet cell types. The gene body of each gene is highlighted.

**Supplementary Figure 2.**
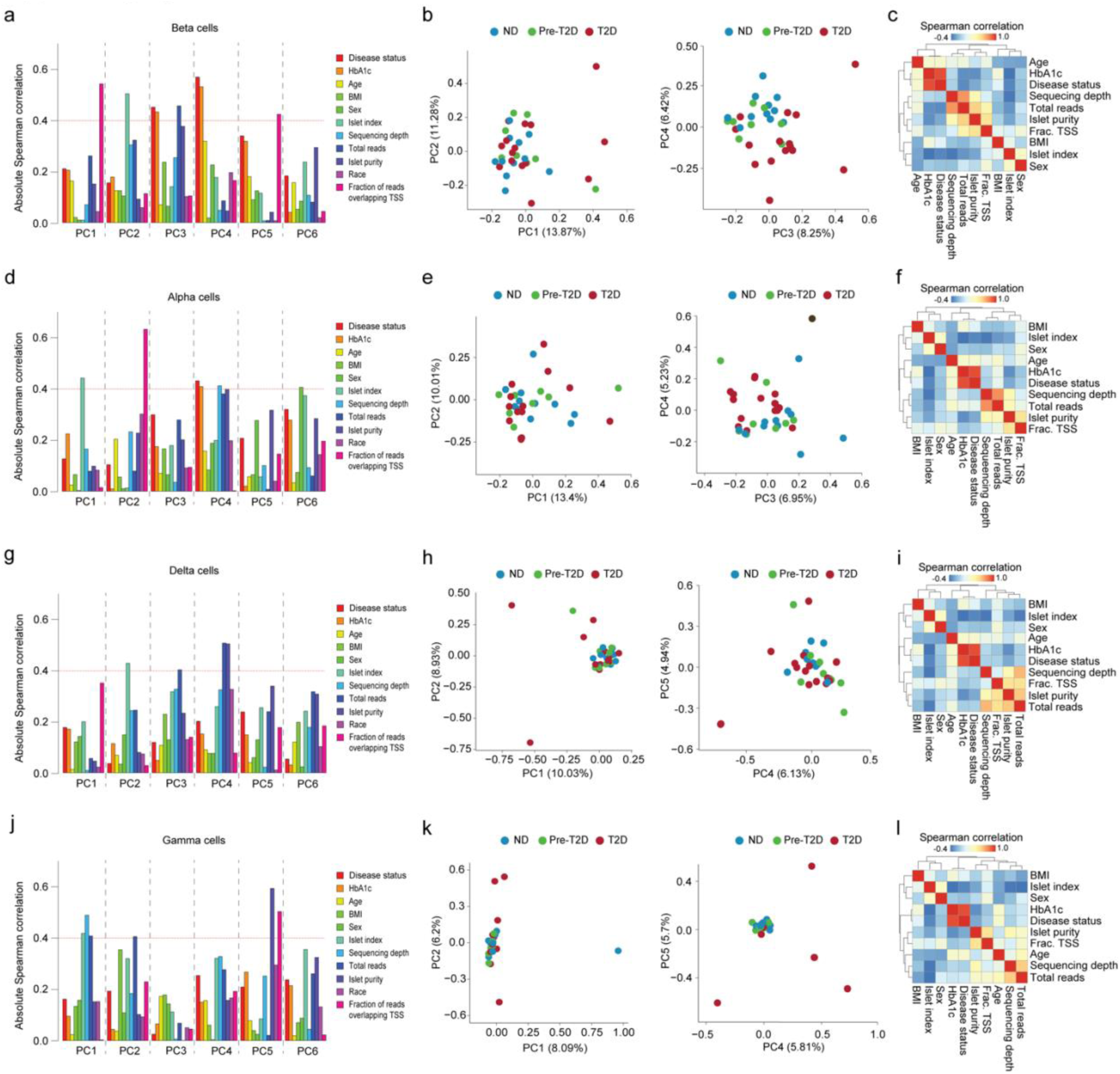
Identification of factors explaining donor variability in snATAC-seq data. **(a,d,g,j)** Absolute Spearman correlation coefficient between the first 6 principle components (PCs) and each biological or technical variable in beta (a), alpha (d), delta (g), and gamma (j) cells. An absolute Spearman correlation threshold of 0.4 was used to identify factors having a high correlation with each PC. **(b,e,h,k)** Principal component analysis (PCA) based on cCREs in beta (b), alpha (e), delta (h), and gamma (k) cells from individual non-diabetic (ND, *n*=11), pre-diabetic (pre-T2D, *n*=8), and type 2 diabetic (T2D, *n*=15) donors which are color-coded by disease status. Each donor in the space is defined by the first two principal components (left) and the two principal components (right) that show highest correlation with disease status. **(c,f,i,l)** Pairwise Spearman correlation coefficients between biological or technical variables in beta (c), alpha (f), delta (i), and gamma (l) cells.

**Supplementary Figure 3.**
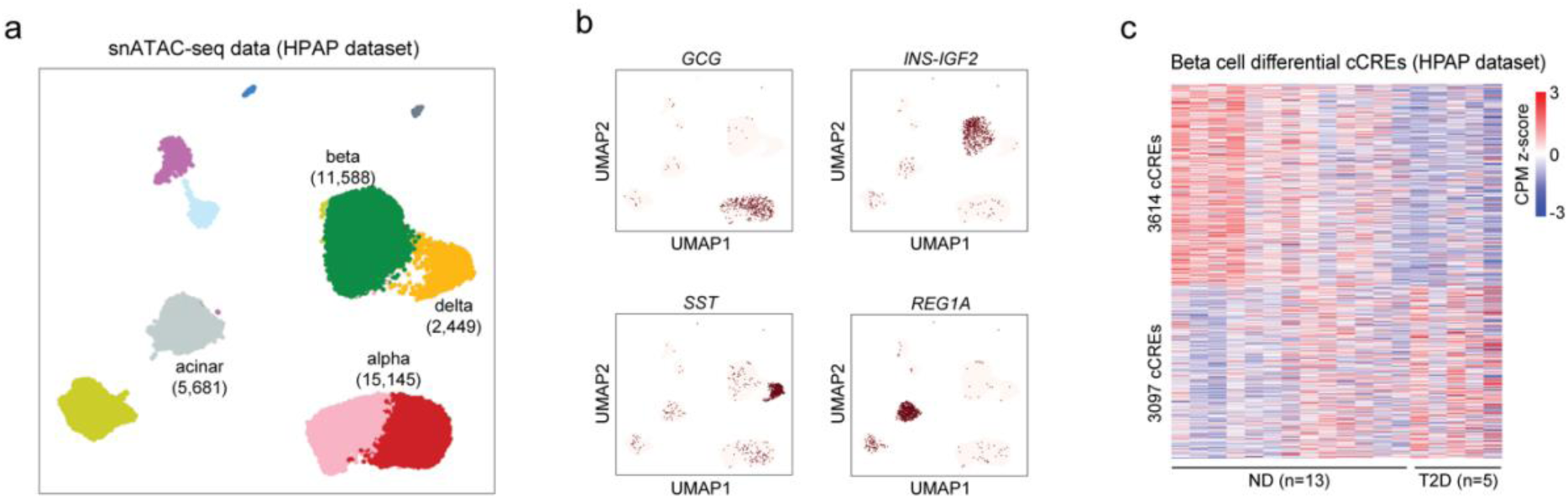
Validation of beta cell T2D-differential cCREs in snATAC-seq data from an independent cohort of donor islets. **(a)** Clustering of chromatin accessibility profiles from HPAP human islet snATAC-seq data (non-diabetic (ND), *n*=13; pre-T2D, *n*=2; T2D, *n*=5). Cells are plotted using the first two UMAP components. Clusters are assigned cell type identities based on promoter accessibility of known marker genes (see Supplementary Figure 3b). The number of cells for each cell type cluster is shown in parentheses. **(b)** Promoter chromatin accessibility in a 5 kb window around TSS for selected endocrine and non-endocrine marker genes for each profiled cell (alpha: *GCG*, beta: *INS-IGF2*, delta: *SST*, acinar: *REG1A*). Total counts normalization and log-transformation were applied. **(c)** Heatmap showing chromatin accessibility at differential cCREs identified in Figure 1e in HPAP snATAC-seq data. Columns represent beta cells from each donor (ND, *n*=13; T2D, *n*=5) and all ND and T2D donors with accessibility of peaks normalized by CPM (counts per million).

**Supplementary Figure 4.**
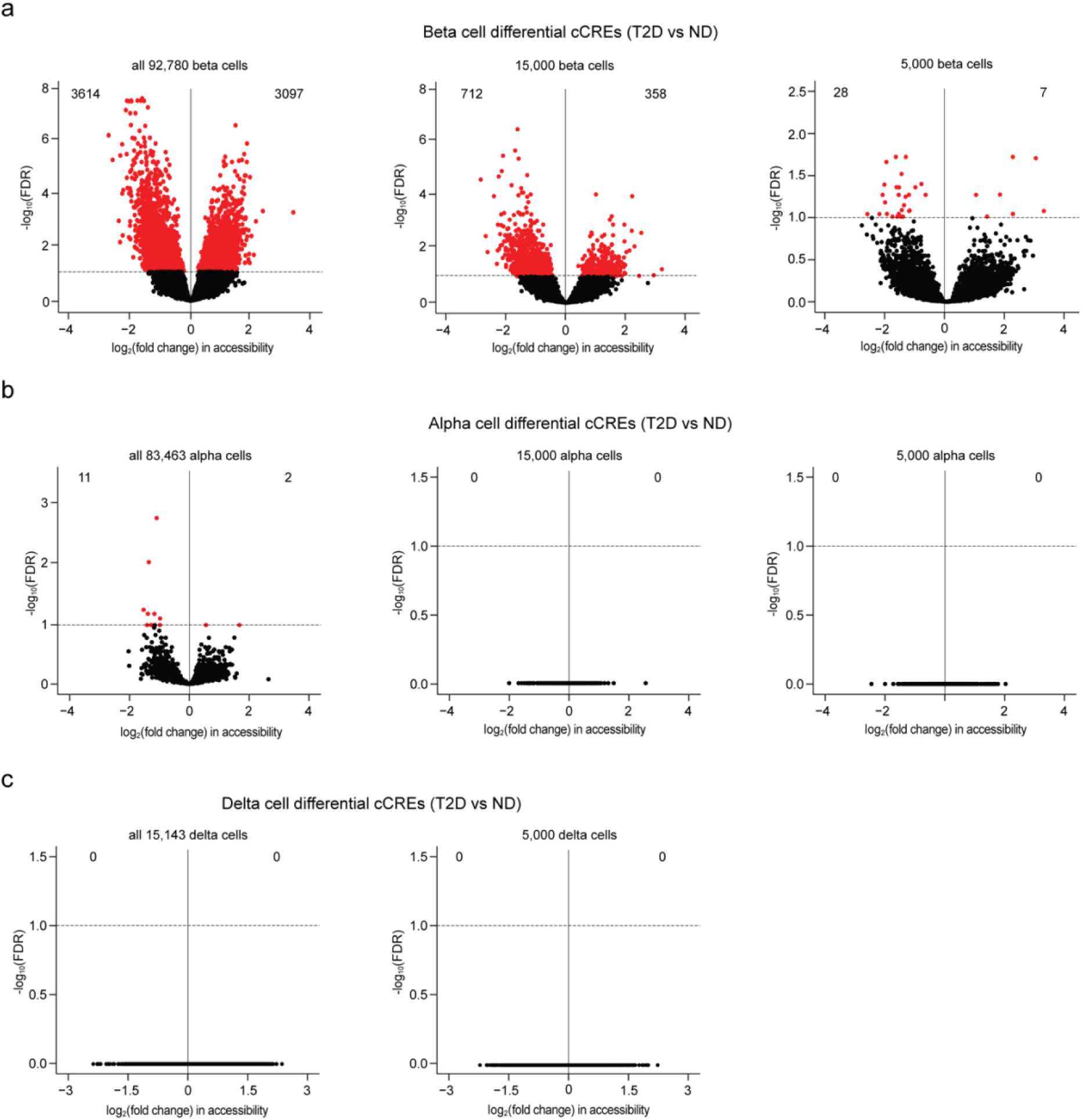
T2D affects chromatin activity more profoundly in beta cells than in other endocrine cell types. **(a)** Volcano plot showing differential cCREs in beta cells between type 2 diabetic (T2D) and non-diabetic (ND) donors. Panels show all beta cells (left), beta cells down-sampled to 15,000 (middle), and 5,000 cells (right). Each dot represents a cCRE. cCREs with FDR < .1 after Benjamini-Hochberg correction (red dots) were considered differentially accessible. **(b)** Volcano plot showing differential cCREs in alpha cells between T2D and ND donors. Panels show all alpha cells (left), alpha cells down-sampled to 15,000 (middle), and 5,000 cells (right). Each dot represents a chromatin accessible cCRE. cCREs with FDR < .1 after Benjamini-Hochberg correction (red dots) were considered differentially accessible. **(c)** Volcano plot showing differential cCREs in delta cells between T2D and ND donors. Panels show all delta cells (left) and delta cells down-sampled to 5,000 cells (right). Each dot represents a chromatin accessible cCRE. cCREs with FDR < .1 after Benjamini-Hochberg correction (red dots) were considered differentially accessible.

**Supplementary Figure 5.**
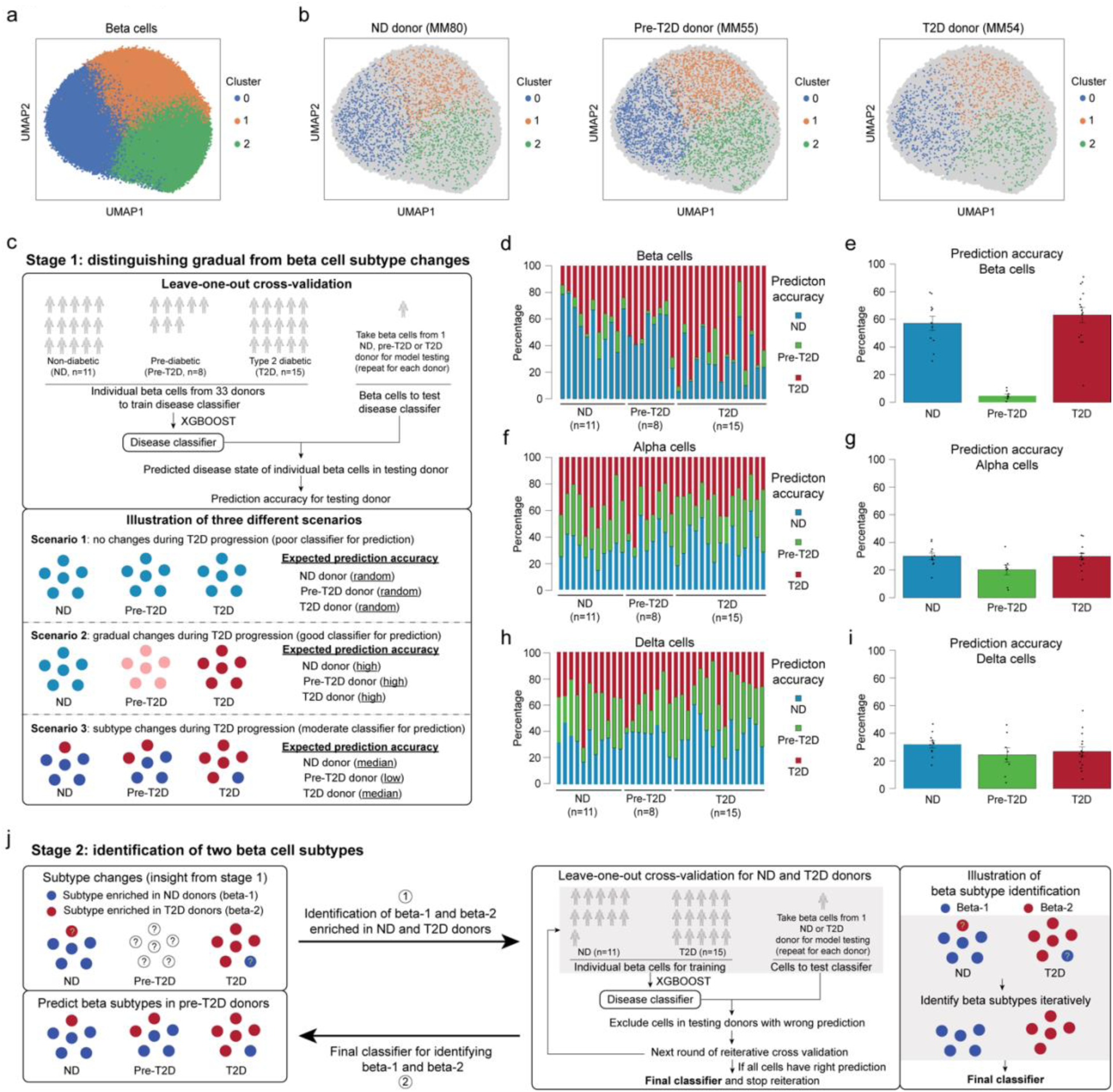
Machine learning undercovers two beta cell subtypes. **(a)** Clustering of chromatin accessibility profiles from 92,780 beta cells from non-diabetic (ND), prediabetic (pre-T2D) and type 2 diabetic (T2D) donor islets using Scanpy (resolution=0.5). Cells are plotted using the first two UMAP components. **(b)** Position of beta cells from representative ND (MM80), pre-T2D (MM55), and T2D (MM54) donors on the UMPA in panel a. **(c)** Illustration of process for distinguishing gradual from subtype changes in beta cells using machine learning. Possible scenarios for cell changes during T2D progression and expected disease state prediction accuracies for each scenario. In the case of no T2D-associated changes, the prediction accuracy for each disease state would be random (scenario 1), gradual cell state changes would be reflected by high prediction accuracy in each disease state (scenario 2), and subtype changes would be reflected by median prediction accuracies (scenario 3, here shown for two cell subtypes). **(d, f, h)** Relative abundance of predicted disease state among beta (d), alpha (f), and delta (h) cells from each donor using XGBOOST. Each column represents cells from one donor. **(e, g, i)** Relative abundance of predicted disease state among beta (e), alpha (g), and delta (i) cells in ND, pre-T2D and T2D donor islets. Data are shown as mean ± S.E.M. (*n* = 11 ND, *n* = 8 pre-T2D, *n* = 15 T2D donors), dots denote data points from individual donors. (**j**) Illustration of process for identifying a classifier capable of distinguishing the two beta cell subtypes.

**Supplementary Figure 6.**
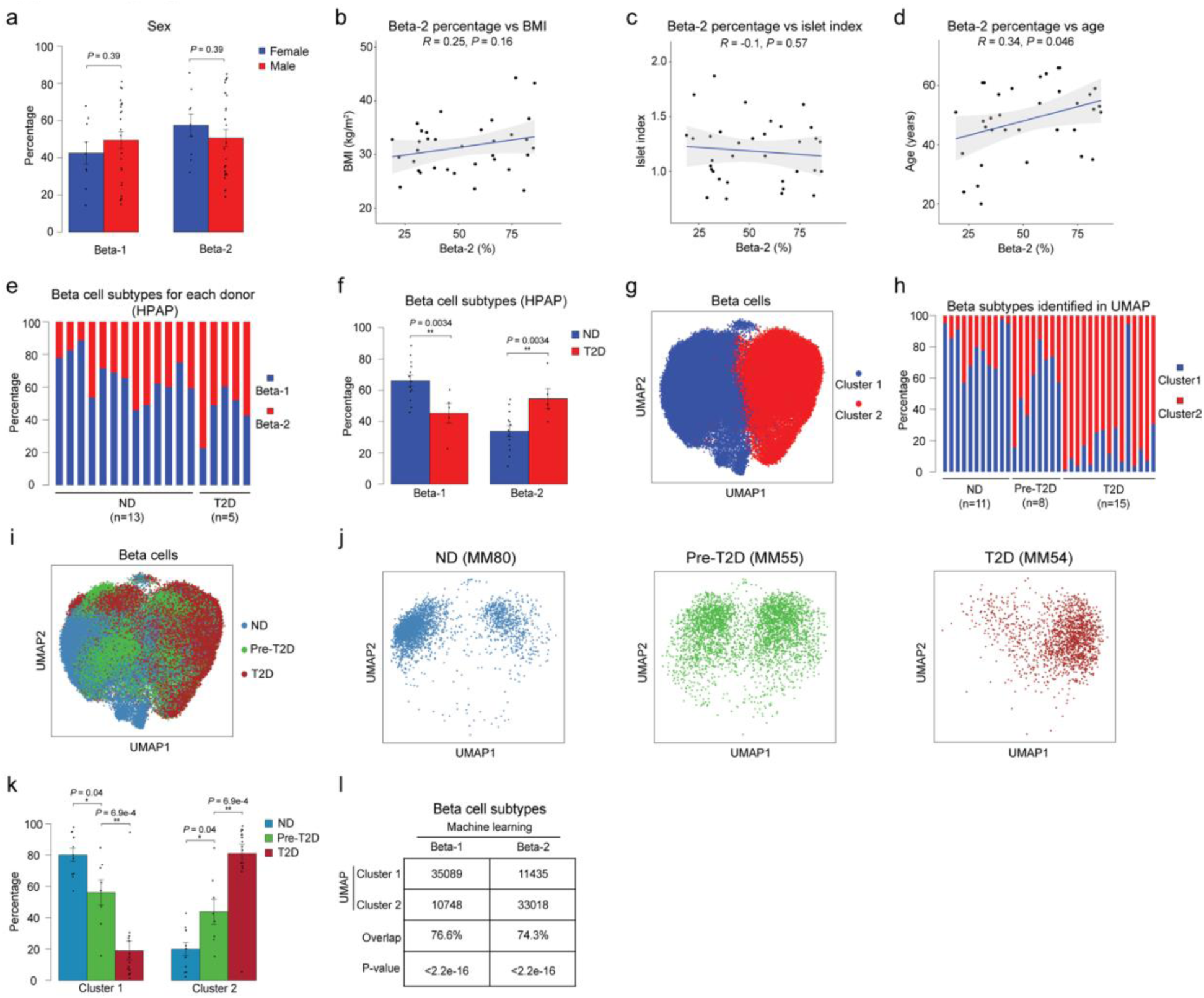
Validation of beta cell subtypes using independent data and computational methods. **(a)** Relative abundance of beta-1 and beta-2 cells in male and female donor islets. Data are shown as mean ± S.E.M. (*n* = 9 females, *n* = 25 males), dots denote data points from individual donors. ANOVA test with age, disease, BMI, and islet index as covariates. **(b)** Pearson correlation between relative abundance of beta-2 cells and BMI across donors (*n* = 11 ND, *n* = 8 pre-T2D, *n* = 15 T2D donors). **(c)** Pearson correlation between relative abundance of beta-2 cells and islet index across donors. **(d)** Pearson correlation between relative abundance of beta-2 cells and age across donors. **(e)** Relative abundance of beta-1 and beta-2 cells in islet snATAC-seq data from an independent cohort (*n* = 13 ND, *n* = 5 T2D donors). Each column represents cells from one donor. **(f)** Relative abundance of each beta cell subtype in ND and T2D donor islets. Data are shown as mean ± S.E.M (*n* = 13 ND, *n* = 5 T2D donors). \*\**P* < .01; ANOVA test with age, sex, and BMI as covariates. **(g)** Clustering of chromatin accessibility profiles from 92,780 beta cells from ND, pre-T2D and T2D donors using beta cell differential cCREs between ND and T2D donors from Figure 1e. Cells are plotted using the first two UMAP components. **(h)** Relative abundance of each beta cell cluster based on UMAP annotation in panel g. Each column represents cells from one donor. **(i)** Position of beta cells from ND, pre-T2D and T2D donors on the UMPA in panel g. **(j)** Position of beta cells from representative ND (MM80), pre-T2D (MM55) and T2D (MM54) donors on the UMPA in panel g. **(k)** Relative abundance of each beta cell cluster in ND, pre-T2D and T2D donor islets. Data are shown as mean ± S.E.M. (*n* = 11 ND, *n* = 8 pre-T2D, *n* = 15 T2D donors). \*\**P* < .01, \**P* < .05; ANOVA test with age, sex, BMI, and islet index as covariates. **(l)** Overlap between beta cell subtypes identified using machine learning and beta cell clusters from UMPA in panel g. The overlap is 76.6% between cluster 1 and beta-1 and 74.3% between cluster 2 and beta-2. *P* < 2.2e-16 (Binominal test).

**Supplementary Figure 7.**
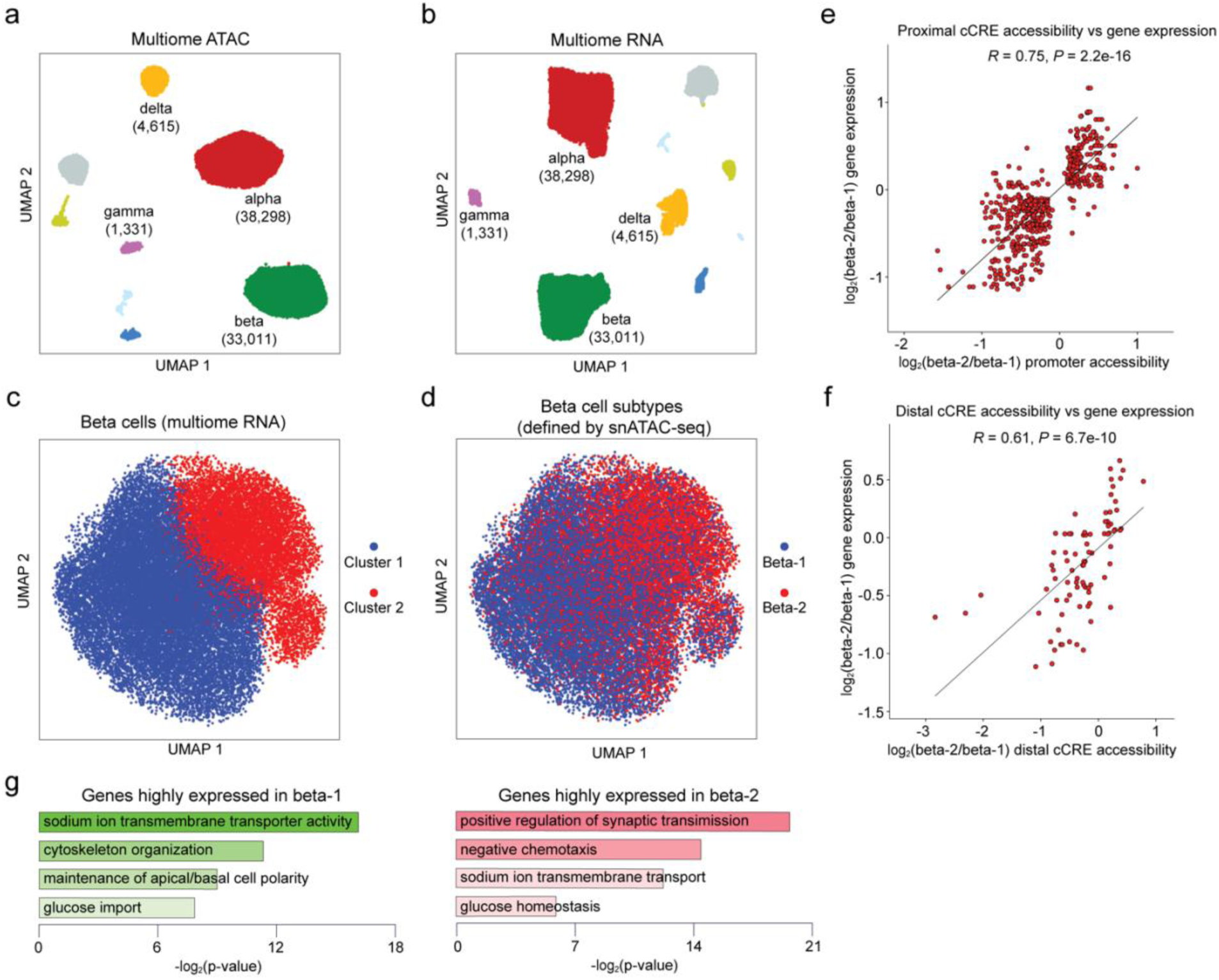
Validation and characterization of beta cell subtypes using multiome data. **(a)** Clustering of chromatin accessibility profiles of cells from multiome data (*n* = 6 ND, *n* = 8 pre-T2D, *n* = 6 T2D). Cells are plotted using the first two UMAP components. Clusters are assigned cell type identities based on promoter accessibility of known marker genes (alpha: *GCG*, beta: *INS-IGF2*, delta: *SST*, gamma: *PPY*). The number of cells for each cell type cluster is shown in parentheses. **(b)** Clustering of gene expression profiles of cells from multiome data (*n* = 6 ND, *n* = 8 pre-T2D, *n* = 6 T2D). Cells are plotted using the first two UMAP components. Clusters are assigned cell type identities based on expression levels of known marker genes (alpha: *GCG*, beta: *INS*, delta: *SST*, gamma: *PPY*). The number of cells for each cell type cluster is shown in parentheses. **(c)** Clustering of gene expression profiles of beta cells from multiome data using genes linked to differential proximal (within ± 5kb of a TSS in GENCODE V19) and distal (based on potential distal cCRE-promoter connections inferred from cicero, see Methods) cCREs between ND and T2D beta cells from Figure 1e. Cells are plotted using the first two UMAP components. **(d)** Plots of beta cell subtypes predicted from chromatin accessibility profiles of beta cells from multiome data by machine learning. **(e)** Correlation between changes in proximal cCRE (within ± 5kb of a TSS in GENCODE V19) accessibility and gene expression differences between beta-1 and beta-2 cells for differentially expressed genes from Figure 3b. There are 544 proximal cCREs and target gene pairs in total, of which 511 have consistent changes between proximal cCRE accessibility and gene expression. **(f)** Correlation between changes in distal cCRE (potential distal cCRE-promoter connections inferred from cicero, see Methods) accessibility and gene expression differences between beta-1 and beta-2 cells for differentially expressed genes from Figure 3b. There are 85 distal cCREs and target gene pairs in total, of which 72 have consistent changes between distal cCRE accessibility and gene expression. **(g)** Enriched gene ontology terms among genes (see Figure 3b) with higher (proximal or distal) cCRE accessibility and expression in beta-1 compared to beta-2 cells (left) and higher (proximal or distal) cCRE accessibility and expression in beta-2 compared to beta-1 cells (right).

**Supplementary Figure 8.**
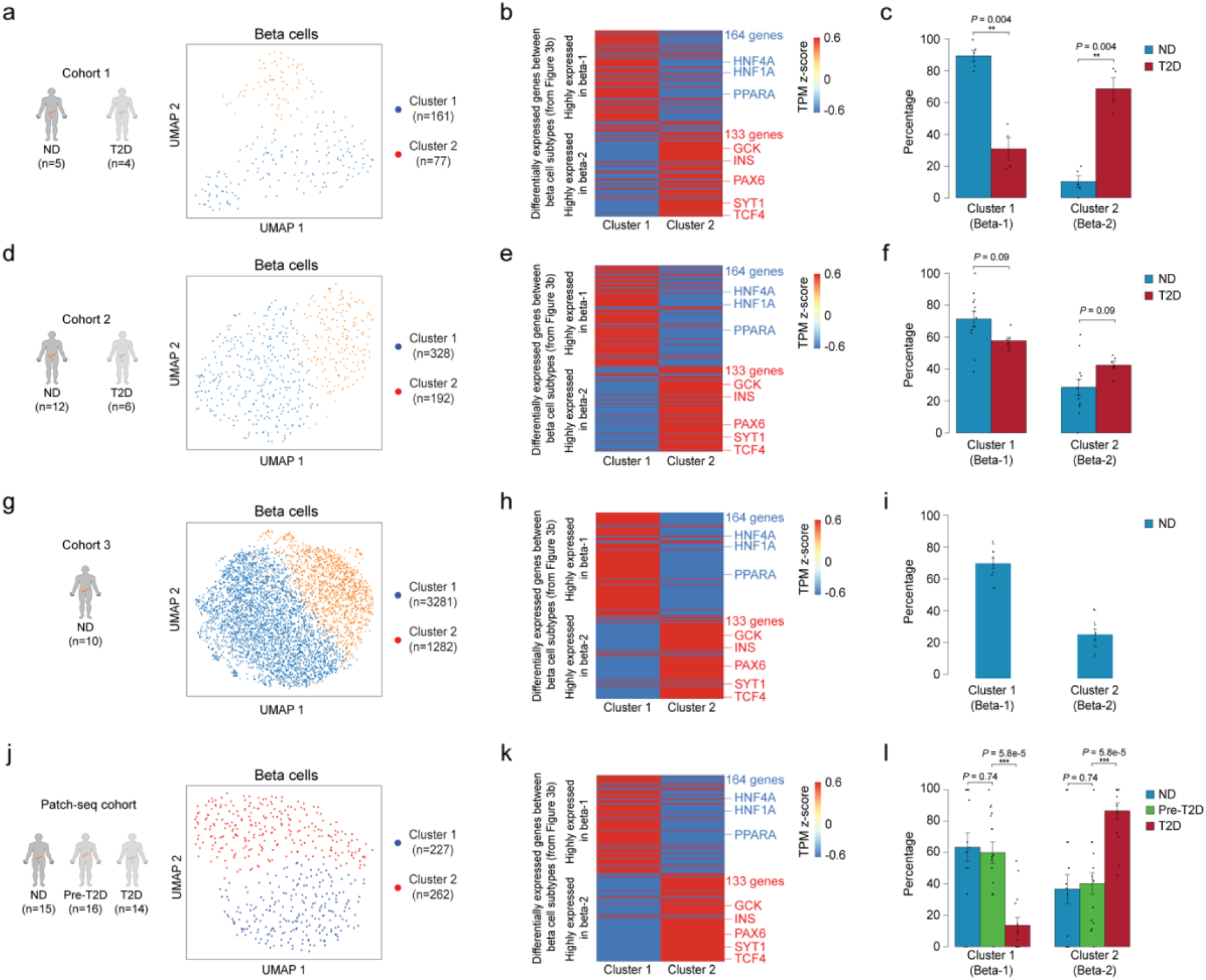
Beta-1 and beta-2 cell classification analysis in scRNA-seq data from independent cohorts. **(a, d, g, j)** Clustering of gene expression profiles of beta cells from cohort 1^5^, cohort 2^12^, cohort 3^22^, and Patch-seq cohort using differentially expressed genes between beta-1 and beta-2 from Figure 3b. Cells are plotted using the first two UMAP components. The number of donors for each cohort and cells for each cell cluster is shown in parentheses. **(b, e, h, k)** Heatmap showing pseudo-bulk expression levels of differentially expressed genes between beta-1 and beta-2 (see Figure 3b) in beta cells from cluster 1 and cluster 2 of cohort 1^5^, cohort 2^12^, cohort 3^22^, and Patch-seq cohort. Expression levels of genes are normalized by TPM (transcripts per million). **(c, f, i, l)** Relative abundance of each beta cell subtype in ND and T2D donor islets in cohort 1^5^, cohort 2^12^, cohort 3^22^, and Patch-seq cohort. Data are shown as mean ± S.E.M., dots denote data points from individual donors. \*\**P* < .01, \*\*\**P* < .001; ANOVA test with age, sex, and BMI as covariates.

**Supplementary Figure 9.**
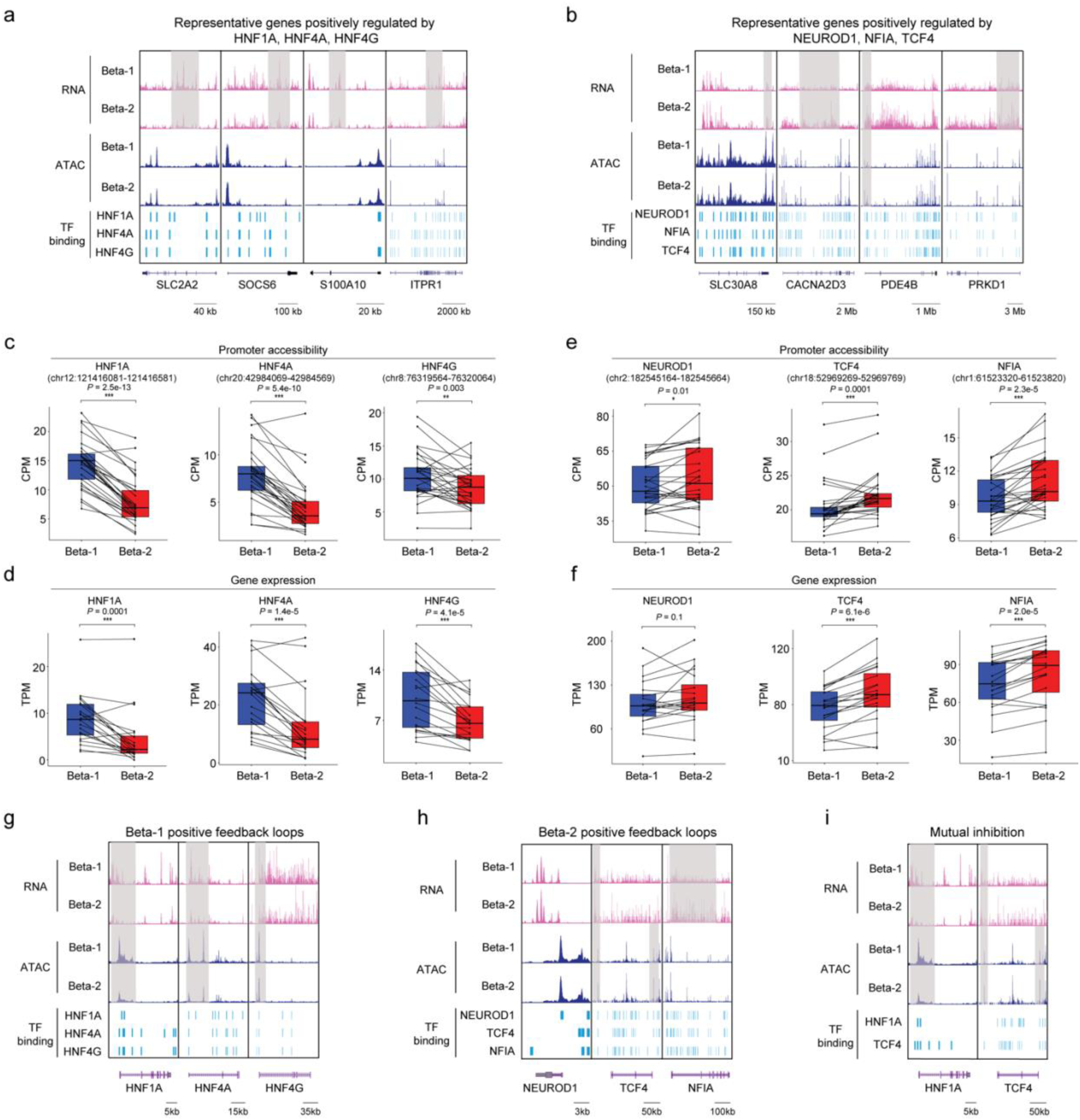
Transcriptional programs distinguishing the two beta cell subtypes. **(a)** Genome browser tracks showing aggregate RNA and ATAC read density at representative genes (*SLC2A2, SOCS6, S100A10, ITPR1*) positively regulated by HNF1A, HNF4A or HNF4G. Differential regions between beta-1 and beta-2 are indicated by grey shaded boxes. Beta cell cCREs with binding sites for HNF1A, HNF4A and HNF4G are shown. All tracks are scaled to uniform 1×10^6^ read depth. **(b)** Genome browser tracks showing aggregate RNA and ATAC read density at representative genes (*SLC30A8, CACNA2D3, PDE4B, PRKD1*) positively regulated by NEUROD1, NFIA or TCF4. Differential regions between beta-1 and beta-2 are indicated by grey shaded boxes. Beta cell cCREs with binding sites for NEUROD1, NFIA and TCF4 are shown. All tracks are scaled to uniform 1×10^6^ read depth. **(c)** Bar plots showing accessibility at *HNF1A*, *HNF4A* and *HNF4G* proximal cCREs in beta-1 and beta-2 cells. Proximal region of *HNF1A* (chr12:121416081-121416581), *HNF4A* (chr20:42984069-42984569), *HNF4G* (chr8:76319564-76320064). Accessibility of peaks is normalized by CPM (counts per million). Paired t-test. **(d)** Bar plots showing expression of *HNF1A*, *HNF4*A and *HNF4G* in beta-1 and beta-2 cells. Gene expression is normalized by TPM (transcripts per million). Paired t-test. **(e)** Bar plots showing accessibility at *NEUROD1*, *NFIA* and *TCF4* proximal cCREs in beta-1 and beta-2 cells. Proximal region of *NEUROD1* (chr2:182545164-182545664), *NFIA* (chr1:61523320-61523820), *TCF4* (chr18:52969269-52969769). Accessibility of peaks is normalized by CPM. Paired t-test. **(f)** Bar plots showing expression of *NEUROD1*, *NFI*A, and *TCF4* in beta-1 and beta-2. Gene expression is normalized by TPM. Paired t-test. **(g)** Genome browser tracks showing aggregate RNA and ATAC read density at *HNF1A, HNF4A* and *HNF4G* in beta-1 and beta-2 cells. Differential regions between beta-1 and beta-2 are indicated by grey shaded boxes. Beta cell cCREs with binding sites for HNF1A, HNF4A and HNF4G are shown. All tracks are scaled to uniform 1×10^6^ read depth. **(h)** Genome browser tracks showing aggregate RNA and ATAC read density at *NEUROD1, NFIA* and *TCF4* in beta-1 and beta-2 cells. Differential regions between beta-1 and beta-2 are indicated by grey shaded boxes. Beta cell cCREs with binding sites for NEUROD1, NFIA and TCF4 are shown. All tracks are scaled to uniform 1×10^6^ read depth. **(i)** Genome browser tracks showing aggregate RNA and ATAC read density at *HNF1A* and *TCF4* in beta-1 and beta-2 cells. Differential regions between beta-1 and beta-2 cells are indicated by grey shaded boxes. Beta cell cCREs with binding sites for HNF1A and TCF4 are shown. All tracks are scaled to uniform 1×10^6^ read depth.

**Supplementary Figure 10.**
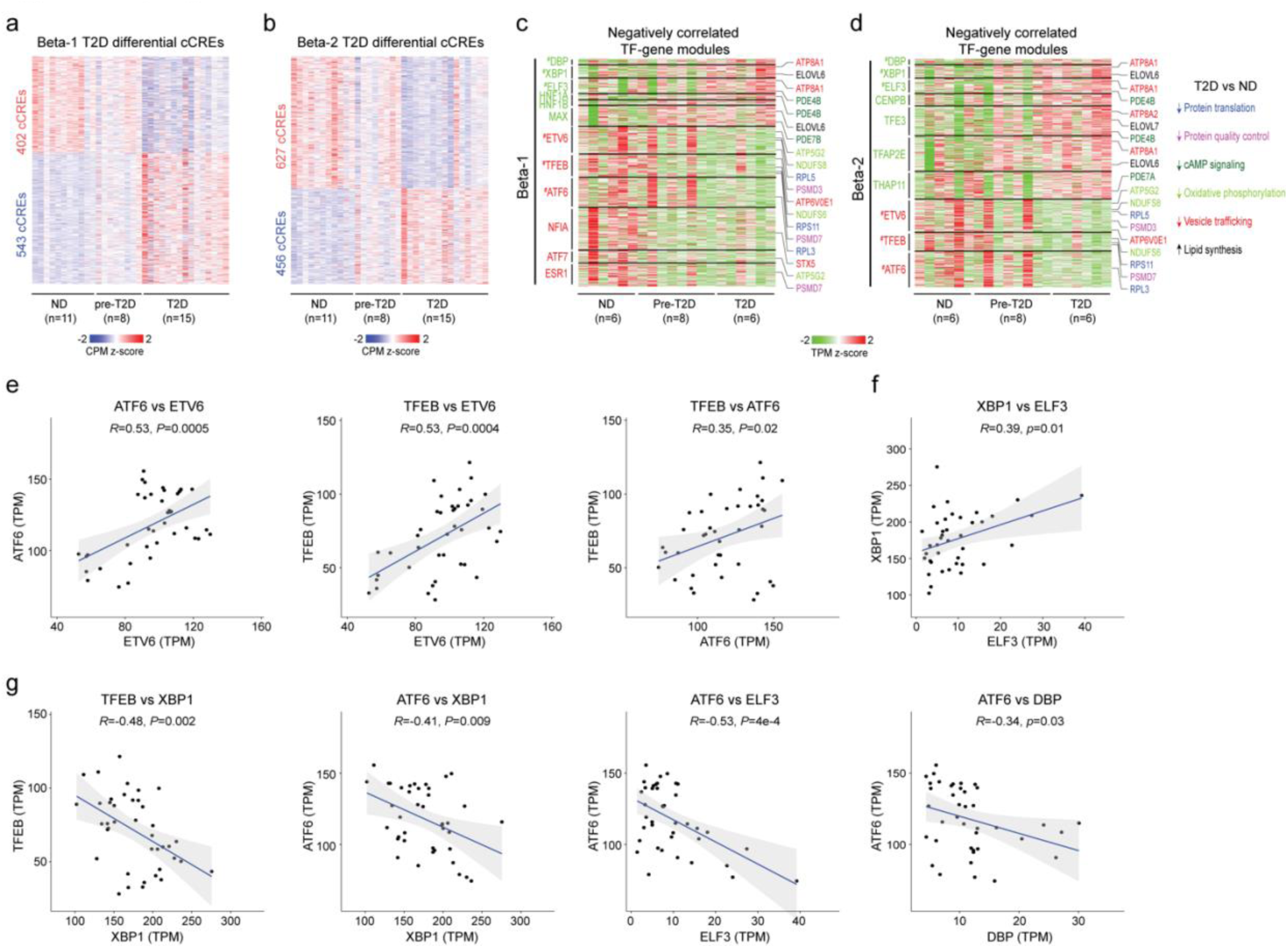
Transcriptional programs changed in both beta cell subtypes in T2D. **(a)** Heatmap showing chromatin accessibility at cCREs with differential accessibility in beta-1 cells from ND and T2D donors. Columns represent beta cells from each donor (ND, *n*=11; pre-diabetic, pre-T2D, *n*=8; T2D, *n*=15) with accessibility of peaks normalized by CPM (counts per million). **(b)** Heatmap showing chromatin accessibility at cCREs with differential accessibility in beta-2 cells from ND and T2D donors. Columns represent beta cells from each donor (ND, *n*=11; pre-diabetic, pre-T2D, *n*=8; T2D, *n*=15) with accessibility of peaks normalized by CPM. **(c)** Heatmap showing expression of genes negatively regulated by TFs (green) with higher activity in ND compared to T2D beta-1 cells (see Methods) and TFs (red) with lower activity in ND compared to T2D beta-1 cells (*n*=6 ND, *n*=8 pre-T2D, *n*=6 T2D donors). Representative target genes of individual TFs are highlighted and classified by biological processes. Gene expression is normalized by TPM (transcripts per million). ^#^ denotes TFs with decreased or increased expression in T2D in both beta-1 and beta-2 cells. **(d)** Heatmap showing expression of genes negatively regulated by TFs (green) with higher activity in ND compared to T2D beta-2 cells (see Methods) and TFs (red) with lower activity in ND compared to T2D beta-2 cells (*n*=6 ND, *n*=8 pre-T2D, *n*=6 T2D donors). Representative target genes of individual TFs are highlighted and classified by biological processes. Gene expression is normalized by TPM (transcripts per million). ^#^ denotes TFs with decreased or increased expression in T2D in both beta-1 and beta-2 cells. **(e,f,g)** Pearson correlation of expression levels between indicated TFs across pseudo-bulk RNA profiles from each beta cell subtype (40 dots in total: 20 donors including *n* = 6 ND, *n* = 8 pre-T2D, *n* = 6 T2D).

**Supplementary Figure 11.**
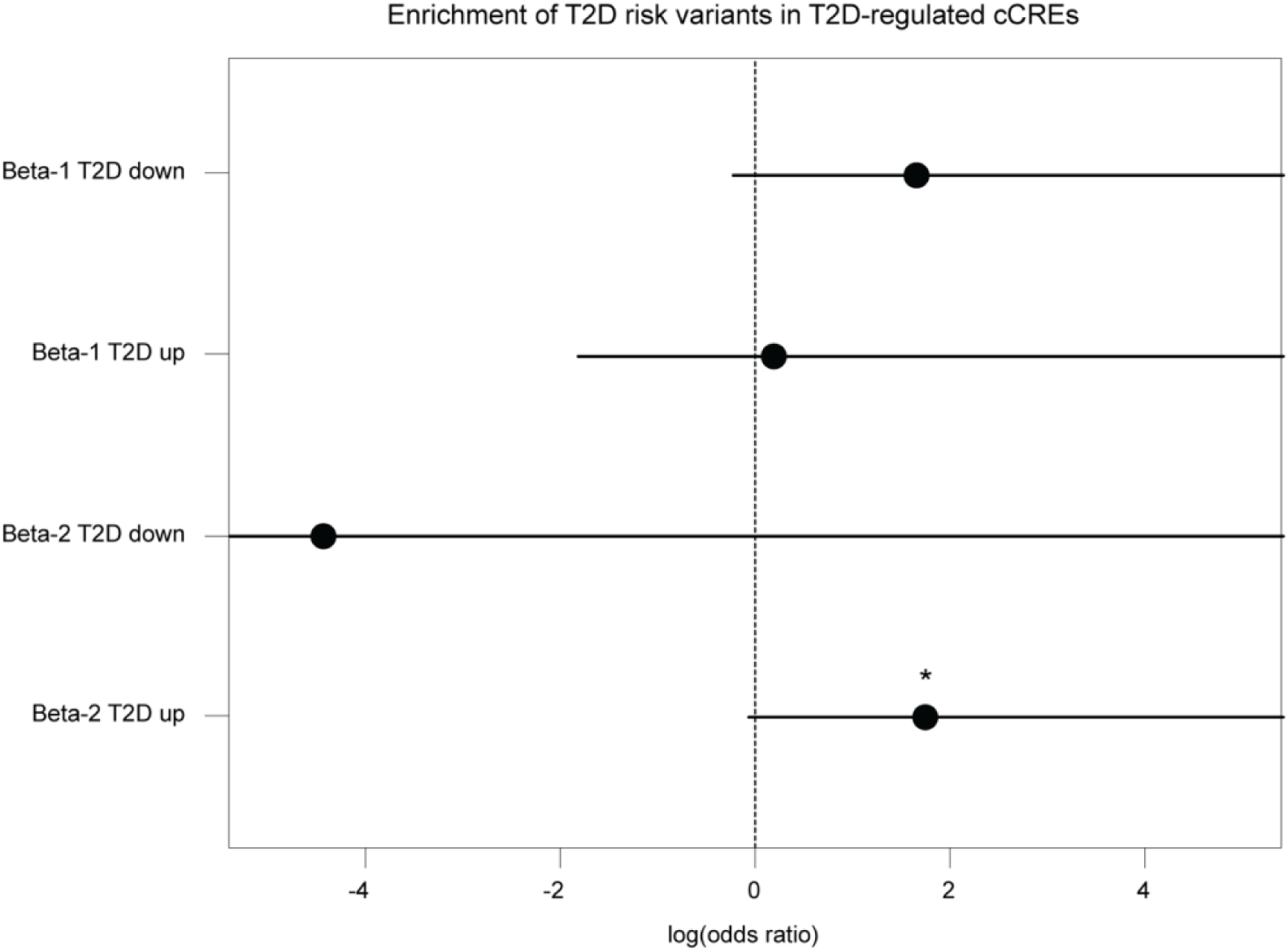
T2D risk variant enrichment for cCREs with T2D-dependent changes in the beta-1 and beta-2 subtype. Enrichment of fine-mapped T2D risk variants for cCREs active in the beta-1 and beta-2 subtype with increased or decreased activity in T2D. Values represent log odds ratios and 95% confidence intervals. * *P* < .05

## References

1. Noguchi, G.M. & Huising, M.O. Integrating the inputs that shape pancreatic islet hormone release. Nat Metab 1, 1189–1201 (2019).

2. Wojtusciszyn, A., Armanet, M., Morel, P., Berney, T. & Bosco, D. Insulin secretion from human beta cells is heterogeneous and dependent on cell-to-cell contacts. Diabetologia 51, 1843–1852 (2008).

3. Dominguez-Gutierrez, G., Xin, Y. & Gromada, J. Heterogeneity of human pancreatic beta-cells. Mol Metab 27S, S7–S14 (2019).

4. Benninger, R.K.P. & Kravets, V. The physiological role of beta-cell heterogeneity in pancreatic islet function. Nat Rev Endocrinol (2021).

5. Segerstolpe, A., et al. Single-Cell Transcriptome Profiling of Human Pancreatic Islets in Health and Type 2 Diabetes. Cell Metab 24, 593–607 (2016).

6. Chiou, J., et al. Single-cell chromatin accessibility identifies pancreatic islet cell type– and state-specific regulatory programs of diabetes risk. Nature Genetics (2021).

7. Dorrell, C., et al. Human islets contain four distinct subtypes of beta cells. Nat Commun 7, 11756 (2016).

8. Cohrs, C.M., et al. Dysfunction of Persisting beta Cells Is a Key Feature of Early Type 2 Diabetes Pathogenesis. Cell Rep 31, 107469 (2020).

9. Chen, C., Cohrs, C.M., Stertmann, J., Bozsak, R. & Speier, S. Human beta cell mass and function in diabetes: Recent advances in knowledge and technologies to understand disease pathogenesis. Mol Metab 6, 943–957 (2017).

10. Fadista, J., et al. Global genomic and transcriptomic analysis of human pancreatic islets reveals novel genes influencing glucose metabolism. Proc Natl Acad Sci U S A 111, 13924–13929 (2014).

11. Wigger, L., et al. Multi-omics profiling of living human pancreatic islet donors reveals heterogeneous beta cell trajectories towards type 2 diabetes. Nat Metab 3, 1017–1031 (2021).

12. Xin, Y., et al. RNA Sequencing of Single Human Islet Cells Reveals Type 2 Diabetes Genes. Cell Metab 24, 608–615 (2016).

13. Lawlor, N., et al. Single-cell transcriptomes identify human islet cell signatures and reveal cell-type-specific expression changes in type 2 diabetes. Genome Res 27, 208–222 (2017).

14. Fang, Z., et al. Single-Cell Heterogeneity Analysis and CRISPR Screen Identify Key beta-Cell-Specific Disease Genes. Cell Rep 26, 3132–3144 e3137 (2019).

15. Wang, Y.J. & Kaestner, K.H. Single-Cell RNA-Seq of the Pancreatic Islets--a Promise Not yet Fulfilled? Cell Metab 29, 539–544 (2019).

16. Camunas-Soler, J., et al. Patch-Seq Links Single-Cell Transcriptomes to Human Islet Dysfunction in Diabetes. Cell Metab 31, 1017–1031 e1014 (2020).

17. Dai, X.Q., et al. Heterogenous impairment of alpha cell function in type 2 diabetes is linked to cell maturation state. Cell Metab 34, 256–268 e255 (2022).

18. Kahn, S.E., Cooper, M.E. & Del Prato, S. Pathophysiology and treatment of type 2 diabetes: perspectives on the past, present, and future. Lancet 383, 1068–1083 (2014).

19. Love, M.I., Huber, W. & Anders, S. Moderated estimation of fold change and dispersion for RNA-seq data with DESeq2. Genome Biol 15, 550 (2014).

20. Chen, T. & Guestrin, C. XGBoost: A Scalable Tree Boosting System. in Proceedings of the 22nd ACM SIGKDD International Conference on Knowledge Discovery and Data Mining 785–794 (2016).

21. M Sander, A.N., J Kalamaras, H C Ee, G R Martin, and M S German. Genetic analysis reveals that PAX6 is required for normal transcription of pancreatic hormone genes and islet development. Genes & development 11, 1662–1673 (1997).

22. Xin, Y., et al. Pseudotime Ordering of Single Human beta-Cells Reveals States of Insulin Production and Unfolded Protein Response. Diabetes 67, 1783–1794 (2018).

23. Efron, B. & Tibshirani, R. On testing the significance of sets of genes. The Annals of Applied Statistics 1(2007).

24. Cahan, P., et al. CellNet: network biology applied to stem cell engineering. Cell 158, 903–915 (2014).

25. Wang, G., et al. A tumorigenic index for quantitative analysis of liver cancer initiation and progression. Proc Natl Acad Sci U S A (2019).

26. Sansbury, F.H., et al. SLC2A2 mutations can cause neonatal diabetes, suggesting GLUT2 may have a role in human insulin secretion. Diabetologia 55, 2381–2385 (2012).

27. Vlacich, G., Nawijn, M.C., Webb, G.C. & Steiner, D.F. Pim3 negatively regulates glucose-stimulated insulin secretion. Islets 2, 308–317 (2010).

28. Stancill, J.S., et al. Chronic beta-Cell Depolarization Impairs beta-Cell Identity by Disrupting a Network of Ca(2+)-Regulated Genes. Diabetes 66, 2175–2187 (2017).

29. Ye, R., et al. Inositol 1,4,5-trisphosphate receptor 1 mutation perturbs glucose homeostasis and enhances susceptibility to diet-induced diabetes. J Endocrinol 210, 209–217 (2011).

30. Martina, J.A., Diab, H.I., Brady, O.A. & Puertollano, R. TFEB and TFE3 are novel components of the integrated stress response. EMBO J 35, 479–495 (2016).

31. Ohta, Y., et al. Clock Gene Dysregulation Induced by Chronic ER Stress Disrupts beta-cell Function. EBioMedicine 18, 146–156 (2017).

32. Eizirik, D.L., Pasquali, L. & Cnop, M. Pancreatic beta-cells in type 1 and type 2 diabetes mellitus: different pathways to failure. Nat Rev Endocrinol 16, 349–362 (2020).

33. Lytrivi, M., Castell, A.L., Poitout, V. & Cnop, M. Recent Insights Into Mechanisms of beta-Cell Lipo- and Glucolipotoxicity in Type 2 Diabetes. J Mol Biol 432, 1514–1534 (2020).

34. Pratt, E.P.S., Harvey, K.E., Salyer, A.E. & Hockerman, G.H. Regulation of cAMP accumulation and activity by distinct phosphodiesterase subtypes in INS-1 cells and human pancreatic beta-cells. PLoS One 14, e0215188 (2019).

35. Bryan, J., et al. ABCC8 and ABCC9: ABC transporters that regulate K+ channels. Pflugers Arch 453, 703–718 (2007).

36. Yang, Y., et al. The phosphatidylserine flippase beta-subunit Tmem30a is essential for normal insulin maturation and secretion. Mol Ther 29, 2854–2872 (2021).

37. Palu, R.A.S. & Chow, C.Y. Baldspot/ELOVL6 is a conserved modifier of disease and the ER stress response. PLoS Genet 14, e1007557 (2018).

38. Tang, N., et al. Ablation of Elovl6 protects pancreatic islets from high-fat diet-induced impairment of insulin secretion. Biochem Biophys Res Commun 450, 318–323 (2014).

39. Gaulton, K.J. Mechanisms of type 2 diabetes risk loci. Current diabetes reports 17, 1–10 (2017).

40. Nkonge, K.M., Nkonge, D.K. & Nkonge, T.N. The epidemiology, molecular pathogenesis, diagnosis, and treatment of maturity-onset diabetes of the young (MODY). Clin Diabetes Endocrinol 6, 20 (2020).

41. Alonso, L., et al. TIGER: The gene expression regulatory variation landscape of human pancreatic islets. Cell Rep 37, 109807 (2021).

42. Chiou, J., et al. Single-cell chromatin accessibility identifies pancreatic islet cell type- and state-specific regulatory programs of diabetes risk. Nat Genet 53, 455–466 (2021).

43. Tsuchiya, Y., et al. IRE1-XBP1 pathway regulates oxidative proinsulin folding in pancreatic beta cells. J Cell Biol 217, 1287–1301 (2018).

44. Seo, H.Y., et al. Endoplasmic reticulum stress-induced activation of activating transcription factor 6 decreases insulin gene expression via up-regulation of orphan nuclear receptor small heterodimer partner. Endocrinology 149, 3832–3841 (2008).

45. Kirkpatrick, C.L., et al. Hepatic nuclear factor 1alpha (HNF1alpha) dysfunction down-regulates X-box-binding protein 1 (XBP1) and sensitizes beta-cells to endoplasmic reticulum stress. J Biol Chem 286, 32300–32312 (2011).

46. Szabat, M., et al. Reduced Insulin Production Relieves Endoplasmic Reticulum Stress and Induces beta Cell Proliferation. Cell Metab 23, 179–193 (2016).

47. Li, H., et al. The Sequence Alignment/Map format and SAMtools. Bioinformatics 25, 2078–2079 (2009).

48. Lareau, C.A., Ma, S., Duarte, F.M. & Buenrostro, J.D. Inference and effects of barcode multiplets in droplet-based single-cell assays. Nat Commun 11, 866 (2020).

49. Wolf, F.A., Angerer, P. & Theis, F.J. SCANPY: large-scale single-cell gene expression data analysis. Genome Biol 19, 15 (2018).

50. Amemiya, H.M., Kundaje, A. & Boyle, A.P. The ENCODE Blacklist: Identification of Problematic Regions of the Genome. Sci Rep 9, 9354 (2019).

51. Consortium, E.P. An integrated encyclopedia of DNA elements in the human genome. Nature 489, 57–74 (2012).

52. Traag, V.A., Waltman, L. & van Eck, N.J. From Louvain to Leiden: guaranteeing well-connected communities. Sci Rep 9, 5233 (2019).

53. Korsunsky, I., et al. Fast, sensitive and accurate integration of single-cell data with Harmony. Nat Methods 16, 1289–1296 (2019).

54. Satpathy, A.T., et al. Massively parallel single-cell chromatin landscapes of human immune cell development and intratumoral T cell exhaustion. Nat Biotechnol 37, 925–936 (2019).

55. Pliner, H.A., et al. Cicero Predicts cis-Regulatory DNA Interactions from Single-Cell Chromatin Accessibility Data. Mol Cell 71, 858–871 e858 (2018).

56. Ji, X., Li, W., Song, J., Wei, L. & Liu, X.S. CEAS: cis-regulatory element annotation system. Nucleic Acids Res 34, W551–554 (2006).

57. Schep, A.N., Wu, B., Buenrostro, J.D. & Greenleaf, W.J. chromVAR: inferring transcription-factor-associated accessibility from single-cell epigenomic data. Nat Methods 14, 975–978 (2017).

58. Fornes, O., et al. JASPAR 2020: update of the open-access database of transcription factor binding profiles. Nucleic Acids Res 48, D87–D92 (2020).

59. Heinz, S., et al. Simple combinations of lineage-determining transcription factors prime cis-regulatory elements required for macrophage and B cell identities. Mol Cell 38, 576–589 (2010).

60. Kuleshov, M.V., et al. Enrichr: a comprehensive gene set enrichment analysis web server 2016 update. Nucleic Acids Res 44, W90–97 (2016).

61. Taliun, D., et al. Sequencing of 53,831 diverse genomes from the NHLBI TOPMed Program. Nature 590, 290–299 (2021).

62. Das, S., et al. Next-generation genotype imputation service and methods. Nat Genet 48, 1284–1287 (2016).

63. Hinrichs, A.S., et al. The UCSC Genome Browser Database: update 2006. Nucleic Acids Res 34, D590–598 (2006).

64. van de Geijn, B., McVicker, G., Gilad, Y. & Pritchard, J.K. WASP: allele-specific software for robust molecular quantitative trait locus discovery. Nat Methods 12, 1061–1063 (2015).

65. Mahajan, A., et al. Trans-ancestry genetic study of type 2 diabetes highlights the power of diverse populations for discovery and translation. medRxiv 2020.09.22.20198937 (2020).

